# TAM receptors control actomyosin dynamics in osteoclasts via RHOA-COFILIN-MYOSIN II signaling

**DOI:** 10.1101/2024.04.12.589232

**Authors:** Janik Engelmann, Jennifer Zarrer, Max Schmerder, Christian Mess, Deniz Ragipoglu, Kristoffer Riecken, Tal Burstyn-Cohen, Emily J. Alberto, Sourav Ghosh, Carla Rothlin, Klaus Pantel, Carsten Bokemeyer, Eric Hesse, Hanna Taipaleenmäki, Sonja Loges, Isabel Ben-Batalla

**Author notes:** Contributed equally (Sonja Loges, Isabel Ben-Batalla). Corresponding authors: J.E.; I.B.B.; S.L.

## Abstract

The TAM family of receptor tyrosine kinases were recently identified to regulate bone homeostasis by controlling osteoblasts and bone formation. Despite extensive knowledge of TAM receptor function in the mononuclear phagocyte system, the role of TAM receptors in osteoclasts remains largely unknown. Here, we identify a physiological regulatory system including MERTK and TYRO3 in osteoclasts controlling RHOA-ROCK-COFILIN/Myosin II signaling, thereby antagonistically regulating osteoclast-mediated bone remodeling to maintain bone homeostasis. Myeloid-specific *lysozyme M*-mediated deletion of *Mertk* led to increased bone mass in mice. In early stages of RANKL-induced osteoclast differentiation MERTK promotes amoeboid migration mode in osteoclast precursor cells by inducing RHOA-COFILIN-MLC2 pathway leading to increased osteoclast formation. In contrast, TYRO3 inhibits RHOA-ROCK signaling in osteoclast precursor cells thereby inhibiting these processes. Furthermore, we unraveled an inhibitory role of MERTK as well as TYRO3 in osteoclast differentiation and function. In line with this, mice with *cathepsin K*-mediated deletion of *Mertk* and *Tyro3* exhibited an osteoporotic bone phenotype. We found that osteoclast precursor cell morphology dictates its fusion capacity and identified MERTK as a negative regulator of osteoclast fusion. In multinucleated cells, deletion of *Mertk* inhibits actin ring formation by mediating central actomyosin contraction and inactivation of COFILIN. In contrast, spatially well-ordered RHOA activation at adhesion structures, induced by loss of *Tyro3,* improves osteoclast biomechanotransduction to ameliorate podosome belt formation and enhance osteoclast function. By using a syngeneic breast cancer bone metastasis osteolysis models we identified TYRO3 as a bone protective receptor for osteolytic bone diseases, whereas MERTK represents a pharmacologic accessible target to inhibit osteoclast formation because its stimulatory effects of osteoclast precursors prevail the inhibitory effects on mature osteoclasts. Next to the recently uncovered role of MERTK as a target for osteoanabolic therapy MERTK may represent a one-drug two-target treatment strategy to increase osteoblast function and reduce osteoclast formation for treatments of bone diseases.

## Introduction

The TAM family of receptor tyrosine kinases, consisting of TYRO3 (BRT, DTK, RSE, SKY, and TIF), AXL (ARK, TYRO7, and UFO), and MERTK (EYK, NYM, and TYRO12) and their cognate ligands growth-arrest-specific gene-6 (GAS6) and protein S (PROS1) maintain adult tissue homeostasis by regulating key molecular signaling pathways and cellular functions including efferocytosis, the process of phagocytic uptake and clearance of apoptotic cells, angiogenesis and taming local or systemic inflammation to inhibit autoimmunity^1–8^.

Recent work demonstrated antagonistic functional roles of MERTK and TYRO3 in osteoblast differentiation and bone formation with MERTK as a potential target for osteoanabolic therapy in osteolytic bone diseases or bone regeneration^9,10^. Additionally, blocking MERTK affected osteoclastogenesis in different cancer bone metastasis mouse models^9^. Nevertheless, its precise role in osteoclasts remained largely unknown. In contrast, it was already demonstrated that TYRO3 promotes osteoclastogenesis and bone resorption *in vitro* and *in vivo*^11–14^. Several studies suggest that TYRO3 is implicated in the pathogenesis of rheumatoid arthritis (RA), a common autoinflammatory disease inducing bone erosions^13,15,16^. Germline deletion of *Tyro3* reduced RA-induced bone erosions in mice^13^. In contrast, another study demonstrated that soluble TYRO3, which can serve as an extracellular trap for PROS1 or GAS6 to inhibit their activity, correlates with RA disease activity score and treatment of osteoclast cultures with soluble TYRO3 led to increased osteoclast differentiation suggesting a negative regulatory function of TAM receptors in osteoclastogenesis^16^.

Osteoclasts derive from hematopoietic stem cell precursors and are the only cell type able to resorb bone^17^. In hematopoietic cells, TAM receptors were found to govern the development and maturation of natural killer cells, erythrocytes, dendritic cells/Langerhans cells and macrophages^18^. Osteoclasts are controlled by two major cytokines, M-CSF and RANKL, which initiate differentiation by activation of TRAF6/NF𝜅B and MAPK signaling to induce transcription factors such as NFATC1 and MITF, AP1 or CREB^19^. The differentiation of mononuclear precursor cells into multinuclear mature osteoclasts is dependent on transformative cell morphogenesis to acquire specialized structural and functional features^20^. Early osteoclast differentiation stages depend on proliferative, anti-apoptotic and migratory stimuli^17,21,22^. At the bone surface osteoclast precursors undergo cell-cell fusion into multinucleated osteoclasts, which in a next step experience an extended cytoskeletal reorganization in order to resorb bone^23,24^. Non-resorbing osteoclasts are flat and classically display podosomes, complex multiprotein F-actin-based structures, mediating adhesion and migration. Bone resorbing osteoclasts are polarized by forming a basolateral and apical cell membrane. Most importantly, osteoclasts encounter the sealing zone, a thick F-actin ring mediating attachment to bone. It encloses a space between basal cell compartments and the bone surface called “Howship’s lacuna” where vacuolar H^+^ ATPase is present at the ruffled membrane and proteolytic enzymes are secreted. The current model of F-actin organization and podosome patterning in osteoclasts implies that osteoclasts upon maturation form podosomes patterned into podosome clusters. As maturation proceeds, these clusters develop into rings which expand into interconnected belts at the periphery of the cell. In contrast, when osteoclasts are present on bone, they display a thicker and more central and stable F-actin ring^23,24^. One of the central proteins controlling cellular architecture, migration and adhesion is non-muscle myosin II (NMMIIA). Myosin II is a hexameric actin-binding protein that consists of two heavy chains, two essential light chains and two regulatory light chains. Its conformation and function are controlled by phosphorylation of its regulatory light chain. Myosin II is activated by phosphorylation of the regulatory light chain on Ser19 (p^Ser19^MLC2)^25^.We could recently demonstrate that TAM receptors MERTK and TYRO3 control RHOA-Myosin II signaling in osteoblasts thereby regulating key cellular functions such as migration, differentiation and bone formation. Thus, we investigated the function of MERTK and TYRO3 in osteoclast formation and bone resorption.

In clinical practice, osteoclast targeting agents mostly consist of bisphosphonates and denosumab^26^. Routinely used bisphosphonates such as alendronate, pamidronate and zoledronic acid have a nitrogen group which blocks the activity of farnesyl diphosphate synthase (FDPS) thereby inhibiting the prenylation of small GTPases such as RAS, RHOA and RAC1 which are critical signaling mediators for maintaining the osteoclast cytoskeleton^27^. Denosumab is a human monoclonal antibody against RANKL^26^. Despite its proven effectiveness in several diseases associated with bone loss, including osteoporosis or bone metastases in cancer we are still facing major challenges with decreasing effects over time and rebound of high osteoclast activity after withdrawal^28^. Therefore, finding novel osteoclast-inhibiting targets for combinatory or sequential therapy is an unmet medical need.

In this study, we show for the first time that TAM receptor signaling by MERTK and TYRO3 represents a regulatory system in osteoclastogenesis in a differentiation stage-dependent manner. We found that osteoclast precursor cell motility is enhanced by MERTK and inhibited by TYRO3 by regulation of amoeboid migration mode by RHOA-COFILIN/MLC2 pathway. In contrast, osteoclast maturation and function are negatively regulated by TAM receptors MERTK and TYRO3 by inhibition of osteoclast-specific actomyosin dynamics. Unraveling these mechanisms point to MERTK as a novel target to inhibit osteolysis in breast cancer bone metastasis, whereas TYRO3 is bone protective by inhibiting osteoclastogenesis.

## Results

### High bone mass induced by *lysozyme M*-mediated deletion of *Mertk*

To elucidate the role of MERTK and TYRO3 in osteoclastogenesis, we created myeloid-cell specific deletions of *Mertk* and *Tyro3* using *lysozyme M -cre* (*LysM-cre*) mice generating *LysM-cre^+^Mertk^flox/flox^*and *LysM-cre^+^Tyro3^flox/flox^* conditional knockout mice. Deletion of *Mertk* and *Tyro3* was confirmed by RT-qPCR analysis in *ex vivo* osteoclast cultures (Suppl. Fig. 1a and b).

Microcomputed tomography (μCT) analysis of trabecular bone of tibia metaphysis of female 8-week-old *LysM-cre^+^Mertk^flox/flox^* mice in comparison to littermate controls (*LysM-cre^-^Mertk^flox/flox^*) revealed increased bone volume, trabecular number and trabecular thickness, whereas trabecular separation was slightly decreased (Fig. 1a-e). We performed bone histomorphometry and observed decreased osteoclast numbers in *LysM-cre^+^Mertk^flox/flox^* mice, whereas osteoblast numbers were not affected, demonstrating that *Mertk* deficiency in myeloid cells impedes osteoclast formation *in vivo* (Fig. 1f-h). Analysis of TRAP^+^ mononuclear cells in the bone marrow revealed decreased osteoclast precursor cell numbers in *LysM-cre^+^Mertk^flox/flox^* mice (Fig. 1i). Additionally, we measured increased distance to nearest bone surface, suggesting that MERTK might be involved in osteoclast precursor cell migration (Fig. 1j).

**Figure 1.**
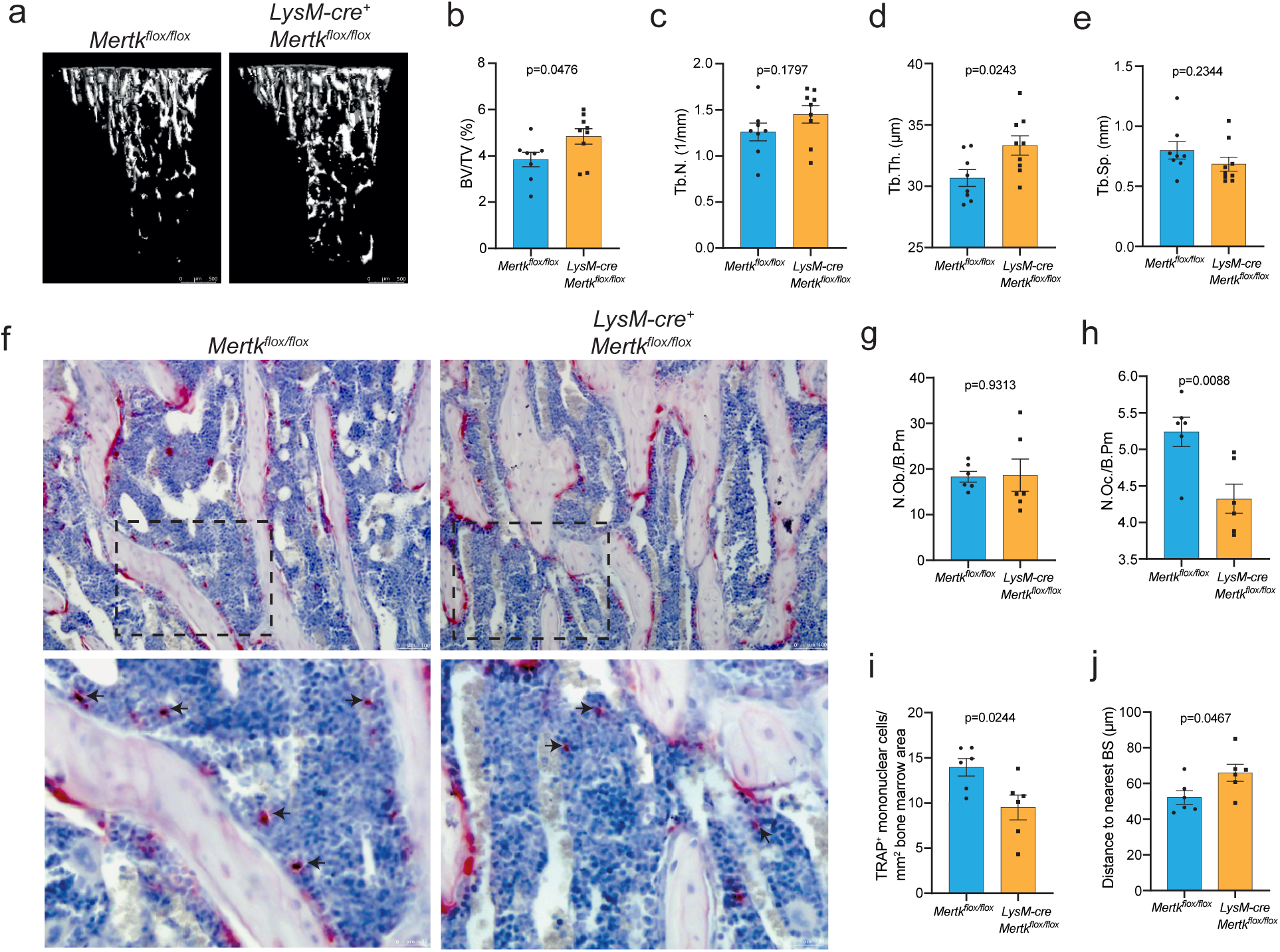
High bone mass induced by *lysozyme M*-mediated deletion of *Mertk*. **a**, Representative images of 3D reconst-ructions of microcomputed tomography (μCT) of trabecular bone of the tibia metaphysis from 8-week-old *Mertk^flox/flox^* and *LysM-cre^+^Mertk^flox/flox^* female mice (longitudinal view of cancellous bone). **b, c, d, e,** Quantification of bone volume (BV/TV) (b), trabecular number (TbN) (c), trabecular thickness (TbTh) (d) and trabecular ^1s00^ eparation (TbSp) (e) of trabecular bone of *Mertk^flox/flox^* and *LysM-cre^+^Mertk^flox/flox^* (n=8/9, mean±SEM, unpaired t-test). **f,** Representative images (upper panel) and magnifications of TRAP/Hematoxylin staining of femur from *Mertk^flox/flox^* and *LysM-cre^+^Mertk^flox/flox^* female mice. **g, h,** Histomorphometric analysis of osteoclast (N.Oc/B.PM) (g) and osteoblast number (N.Ob/B.PM) (h) (n=6/6, mean±SEM, unpaired t-test) **i, j,** Histomorphometric analysis of TRAP^+^ mononuclear cells (i) and osteoclast precursor cell distance to nearest bone surface (black arrows) (j) (n=6/6, mean±SEM, unpaired t-test).

### TAM receptor expression is downregulated by RANKL during osteoclastogenesis

To gain insight into the mechanistic function of MERTK and TYRO3 in osteoclastogenesis, we screened in a first step for differential expression of *Mertk* and *Tyro3* mRNA over the time course of *ex vivo* osteoclast cultures stimulated with M-CSF and RANKL (24 h, 48 h, 72 h and 96 h). Additionally, we measured gene expression of TAM receptor ligands *Pros1* and *Gas6.* RT-qPCR analysis showed that RANKL (50 ng/ml) stimulation downregulates expression of *Mertk* as well *Tyro3*, *Pros1* and *Gas6* and this effect was RANKL-dependent as a low RANKL concentration (10 ng/ml) led to a less pronounced effect (Fig. 2a and b). Therefore, we concluded that MERTK is mainly expressed in osteoclast precursor cells. In concordance with the RT-qPCR data, we detected strong immunoreactivity for MERTK in mononuclear cells (Fig. 2c, white arrows), whereas weaker staining was measured in multinucleated osteoclasts, where MERTK was mainly retained in the nucleus (Fig. 2c, blue and purple arrows).

**Figure 2.**
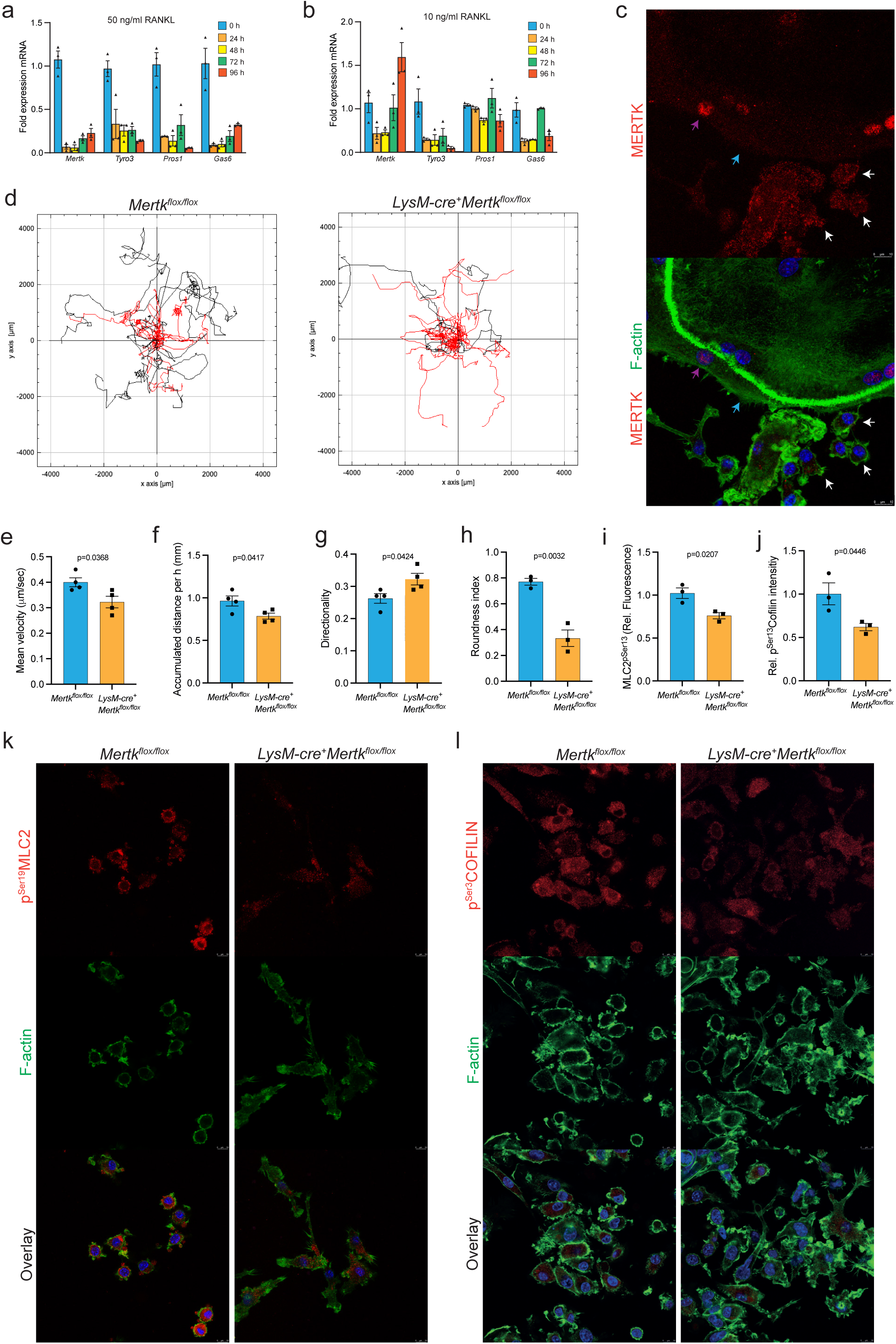
MERTK is mainly expressed in mononuclear osteoclast precursor cells and induces amoeboid migration. **a,b**, mRNA expression levels of TAM receptor family *Mertk*, *Tyro3*, *Axl*, *Gas6*, and *Pros1* in *ex vivo* osteoclast cultures in bone marrow macrophages (0 h) and 24 h, 48 h, 72 h and 96 h after osteoclastogenic induction with M-CSF and the standard RANKL dose (50 ng/ml) (a) and a low*^Loremipsum^* RANKL dose (10 ng/ml) (b). **c,** Representative images of immunofluorescence single cell staining of MERTK in ost*^jjddjips^*e*^um^* oclast cultures. High MERTK signal in the osteoclast nuclei (Purple arrow). Low MERTK expression on osteoclast*^d^* cell surface (Blue arrow). High MERTK expression on mononuclear osteoclast precursor cell surface (White arrow). **d,** Trajectory plots of time-lapse microscopy recordings of single osteoclast precursor cells from from *Mertk^flox/flox^* and *LysM-cre^+-^ Mertk^flox/flox^* mice showing individiual path of migrating cells (Cells with velocity >0,5 μm/sec in black, <0,5μm/sec in red). **e, f, g,** Accompanying analysis of mean velocity (e), accumulated distance per h (f) and directionality (g) (n=4, mean±SEM, unpaired t-test). **h,** Analysis of roundness index of osteoclast precursor cells from *Mertk^flox/flox^* and *LysM-cre^+^Mertk^flox/flox^* mice (n=3 random HPF, mean±SEM, unpaired t-test). **i,** Analysis of p^Ser19^MLC2 immunofluorescence intensity of osteoclast precursor cells from *Mertk^flox/flox^* and *LysM-cre^+^Mertk^flox/flox^* mice (n=3 random HPF, mean±SEM, unpaired t-test). **j,** Analysis of p^Ser3^COFILIN immunofluorescence intensity of osteoclast precursor cells from *Mertk^flox/flox^* and *LysM-cre^+^Mertk^flox/flox^* mice (n=3 random HPF, mean±SEM, unpaired t-test). **k,l,** Representative images of p^Ser19^MLC2 (k) and p^Ser3^COFILIN (l) immunofluorescence staining of osteoclast precursor cells from *Mertk^flox/flox^* and *LysM-cre^+^Mertk^flox/flox^* mice.

### MERTK induces amoeboid migration of osteoclast precursors

Osteoclast precursors are highly mobile cells constantly migrating into and within bone resorption sites^29^. Because of our findings suggesting a role of MERTK in osteoclast precursor migration *in vivo*, we analyzed motility *in vitro* by performing dynamic imaging using time-lapse microscopy. Stimulation with M-CSF and RANKL resulted in high motility of osteoclast precursors with predominance of amoeboid migration mode (Suppl. Video 1). The amoeboid migration has been defined by low adhesion, independence from proteolytic degradation of the extracellular matrix (ECM), and a rounded cell morphology with high actomyosin contractility^30,31^. In contrast, deletion of *Mertk* led to loss of the round, spherical osteoclast precursor cell shape leading to reduced migration velocity and accumulated distance (Suppl. Video 2) (Fig. 2d-f). Interestingly, we observed a more crawling like migratory phenotype induced by loss of *Mertk*, suggesting a shift from amoeboid migration to mesenchymal migration mode, corroborated by increased directionality (Fig. 2g) (Suppl. Video 2). Analysis of roundness index by filamentous actin (F-actin) and actomyosin contractility measured by myosin light chain 2 (MLC2) Ser19 phosphorylation confirmed loss of roundness and contractility induced by deletion of *Mertk* (Fig. 2h, i and k).

Additionally, we stained for Ser3 phosphorylation of COFILIN. Unphosphorylated COFILIN, but not COFILIN phosphorylated at Ser3, can bind actin and promote actin polymerization and depolymerization. Phosphorylated COFILIN inhibits all cofilin–actin interactions and prevents severing of F-actin. Non-phosphorylated COFILIN is found in locomotory and invasive protrusions such as lamellipodia^32^. We detected decreased levels of COFILIN phosphorylated at Ser3 in osteoclast precursors deleted for *Mertk* (Fig. 2j and l). Concomitantly, we observed increased lamellipodia formation in *Mertk*-deficient osteoclast precursors (Suppl. Video 2). In contrast, round spherical control cells exhibited high phosphorylated COFILIN suggesting low actin polymerization (Fig. 2j and l).

### MERTK and TYRO3 antagonistically regulate RHOA during osteoclastogenesis

Next, we investigated upstream regulatory pathways. The major regulator of actomyosin contractility and COFILIN phosphorylation by activation of LIMK is the small GTPase RHOA and its effector Rho associated kinase (ROCK). RHOA act as a molecular switch, triggered by receptor signals between the inactive state as guanosine diphosphate (GDP) and guanosine triphosphate (GTP)-linked action^33^. Pull down assays with Rhotekin-RBD binding GTP-RHOA showed surprisingly that TAM receptor ligand PROS1 dose dependently decreased RHOA activation in control osteoclast precursors in presence and absence of MERTK (Fig. 3A). Therefore, we hypothesized that PROS1 could inhibit RHOA signaling via TYRO3 as there is no other receptor known PROS1 can bind to. Indeed, PROS1 induced strong RHOA activation in absence of TYRO3 which, we suggest, is induced by activation of MERTK (Fig. 3A). In turn, PROS1 increased Ser3 phosphorylation of COFILIN in control osteoclast precursors and in absence of TYRO3, but not MERTK, indicating that PROS1-MERTK axis increases global COFILIN Ser3 phosphorylation via RHOA (Fig. 3a). PROS1-TYRO3 axis reduces global RHOA activity which outweighs the activating effects on RHOA induced by MERTK in baseline.

**Figure 3.**
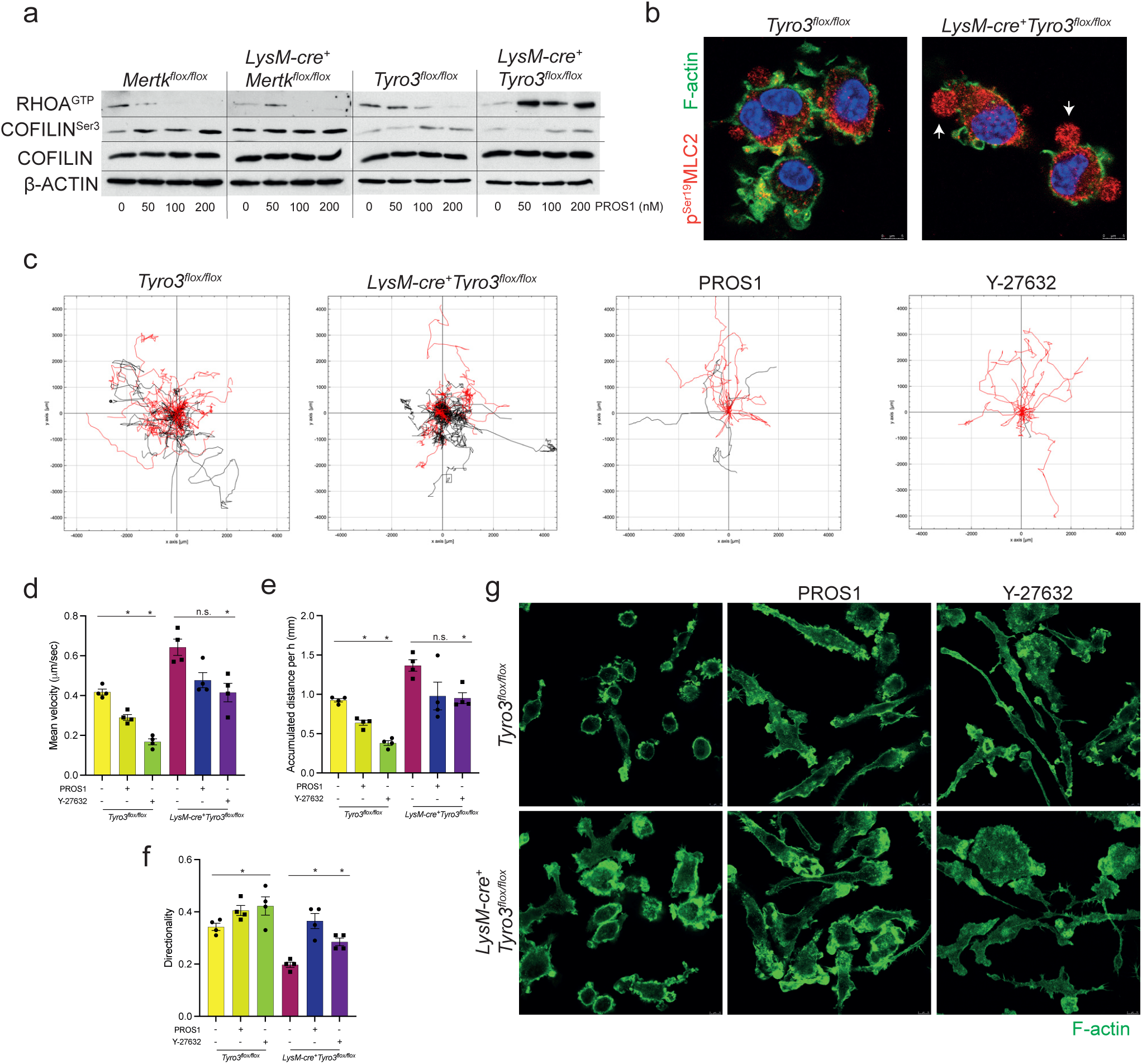
PROS1 inhibits osteoclast precursor amoeboid migration mode by silencing of RHOA-ROCK axis via TYRO3. **a**, Immunoblots of RHOA^GTP^, p^Ser3^COFILIN, COFILIN and β-ACTIN in osteoclast precursor cells from *Mertk^flox/flox^*, *LysM-cre^+^Mertk^flox/flox^* mice, *Tyro3^flox/flox^* and *LysM-cre^+^Tyro3^flox/flox^* mice treated with different concentrations of human plasma purified PROS1. **b,** Representative images of p^Ser13^MLC2 and F-actin immunofluorescence stainings of single osteoclast precursor cells from *Tyro3^flox/flox^* and *LysM-cre^+^Tyro3^flox/flox^* mice. **c,** Trajectory plots of time-lapse microscopy recordings of single osteoclast precursor cells from *Tyro3^flox/flox^* and *LysM-cre^+^Tyro3^flox/flox^* mice and stimulated with PROS1 (200 nM) and the ROCK inhibitor Y-27632 (10 μM) showing individiual path of migrating cells (Cells with velocity >0,5 μm/sec in black, <0,5 μm/sec in red). **d,e,f,** Analysis of mean velocity (d) accumulated distance per h (e) and directionality (f) of single osteoclast precursor cells from *Tyro3^flox/flox^* and *LysM-cre^+^Tyro3^flox/flox^* mice treated with PROS1 (200 nM) and Y-27632 (10 μM) (n=4, mean±SEM, *p<0.05, unpaired t-test). **g,** Representative images of F-actin immunofluorescence stainings of single osteoclast precursor cells from *Tyro3^flox/flox^* and *LysM-cre^+^Tyro3^flox/flox^* mice treated with PROS1 (200 nM) and Y-27632 (10 μM).

### TYRO3 induces mesenchymal migration mode in osteoclast precursors

We next performed immunofluorescence staining of M-CSF and RANKL stimulated osteoclast precursor cells with deletion of *Tyro3.* We observed highly increased formation of multiple membrane protrusions emanating from the cell body in all directions and thus to a loss of polarity (Suppl. Video 3). We observed increased p^Ser19^MLC2 staining intensity, indicating enhanced actomyosin contractility with high numbers of Myosin-induced blebs (Fig. 3b, arrows). Deletion of *Tyro3* led to highly increased velocity and accumulated distance but reduced directionality due to loss of polarity. Treatment of osteoclast precursors with TAM receptor ligand PROS1 led to a shift to mesenchymal migration mode with decreased velocity and accumulated distance but increased directionality (Fig. 3c-f) (Suppl. Video 4). ROCK inhibitor Y-27632 led to similar effects and induced likewise a mesenchymal-like migration mode with decreased velocity and accumulated distance but increased directionality (Fig. 3c-f) (Suppl. Video 5). Treatment of *Tyro3*-deficient osteoclast precursors with Y-27632 could restore velocity, accumulated distance and directionality to control levels, indicating that TYRO3 inhibits ROCK signaling (Fig. 3c-f). In contrast, PROS1 did not affect velocity and accumulated distance, but increased directionality in absence of *Tyro3* confirming that the main receptor for PROS1 in osteoclast precursors is TYRO3, which inhibits RHOA-ROCK signaling (Fig. 3c-f). We performed additional experiments to assess osteoclast precursor cell morphology by F-actin immunofluorescence. We observed again similar effects induced by PROS1 and Y-27632 on precursor cell morphology with induction of a flattened out and spreaded cell phenotype. In absence of *Tyro3*, Y-27632 induced strong effects on cell morphology whereas PROS1 only induced minor changes confirming that PROS1-TYRO3 axis inhibits RHOA-ROCK signaling (Fig. 3g).

### Activation of MERTK and TYRO3 does not lead to increased expression of osteoclast-related genes

To investigate the role of MERTK and TYRO3 in osteoclast differentiation *in vitro,* we performed *ex vivo* osteoclast differentiation cultures obtained from *LysM-cre^+^Mertk^flox/flox^*and *LysM-cre^+^Tyro3^flox/flox^* mice. We observed that deletion of *Mertk* led to decreased, whereas deletion of *Tyro3* led to increased osteoclast numbers. Osteoclast differentiation marker *Nfatc1*, *Acp5*, *Dcstamp* and *Ctsk* were slightly, but inconsistently decreased over the timecourse of differentiation by deletion of Mertk as well as Tyro3 (Suppl. Fig. 2a-f, i-n). Presence of different concentrations of MERTK and TYRO3 ligand PROS1 led to increased osteoclast formation but induced a paradox dose-dependent effect: Lower PROS1 concentrations (50nM) promoted highest numbers of osteoclasts, whereas higher PROS1 levels (100 nM and 200 nM) led to decreased osteoclast formation in comparison to the lower PROS1 dose (Suppl. Fig. 2a and b and 2i and j). Nevertheless, despite its relatively strong effect on osteoclast formation, PROS1 only weakly and inconsistently increased osteoclast differentiation marker *Nfatc1*, *Acp5*, *Dcstamp* and *Ctsk* (Suppl. Fig. 2a-f, i-n). Although total osteoclast number was reduced by deletion of *Mertk*, we observed increased percentage of mature osteoclasts exhibiting a flat morphology with actin ring formation in the cell periphery, suggesting that MERTK may inhibit late osteoclast differentiation and maturation (Suppl. Fig. 2g and h). Likewise, deletion of *Tyro3* led to increased osteoclast maturity (Suppl. Fig. 2o and p). Therefore, we concluded that decreased osteoclast formation induced by deletion of *Mertk in vitro* and *in vivo* is unlikely due to a positive regulation of osteoclast differentiation by MERTK. Moreover, these results suggest a negative regulatory function of MERTK and TYRO3 in late osteoclast differentiation and maturation.

### Osteoporotic bone phenotype induced by *cathepsin K*-mediated deletion of *Mertk* and Tyro3

For investigating late osteoclast differentiation and function, we used *cathepsin K* promotor for osteoclast-specific cre recombinase expression (*Ctsk-cre*)*. Ctsk-cre* mediated gene deletion is mainly targeted in already multinucleated cells, but not in precursor cells^34^. We generated *Ctsk-cre^+^Mertk^flox/flox^*and *Ctsk-cre^+^Tyro3^flox/flox^* mice and performed *ex vivo* osteoclast differentiation assay to confirm deletion of *Mertk* and *Tyro3*, respectively by RT-qPCR (Suppl. Fig. 1c and d). As hypothesized, we observed decreased bone volume in *Ctsk-cre^+^Mertk^flox/flox^* mice (Fig. 4a and b). Likewise, trabecular number and trabecular thickness were decreased, whereas trabecular separation was increased (Fig. 4c-f). Histomorphometric analysis revealed that osteoclast numbers were increased, whereas osteoblast numbers were not affected (Fig. 4b, g and h), indicating that loss of *Mertk* promotes late osteoclast differentiation and accelerates osteoclastic bone resorption. Analysis of bone volume in *Ctsk-cre^+^Tyro3^flox/flox^*mice revealed similarly decreased bone volume, trabecular number and trabecular thickness, whereas trabecular separation was increased (Fig. 4a and i-l). Osteoclast numbers were increased, whereas osteoblast numbers were not affected by deletion of *Tyro3* in osteoclasts (Fig. 4b, m and n). These results demonstrate that loss of *Mertk* as well as *Tyro3* promotes late osteoclast differentiation and accelerates osteoclastic bone resorption resulting in an osteoporotic bone phenotype.

**Figure 4.**
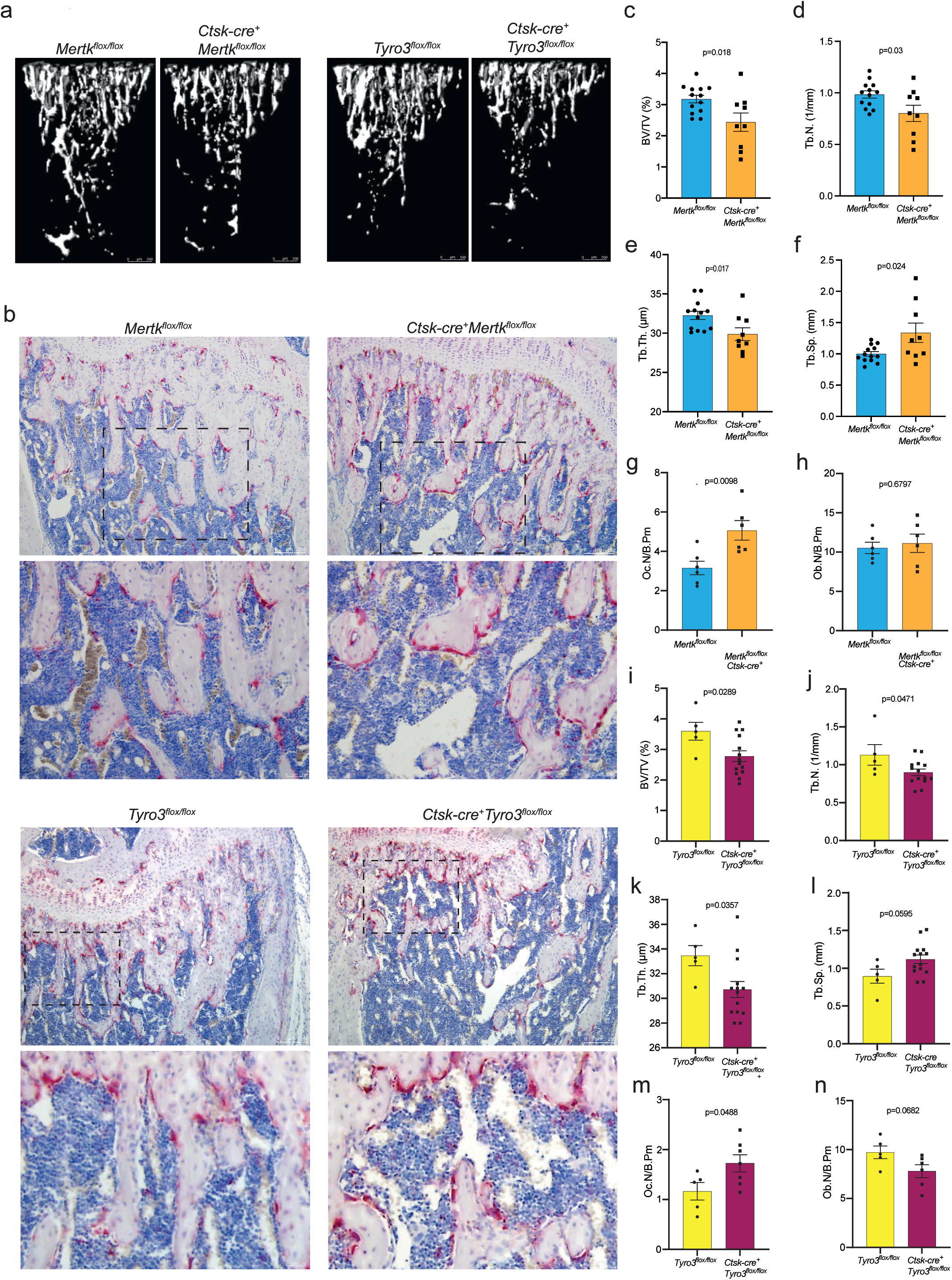
Osteoporotic bone phenotype induced by cathepsin K-mediated deletion of *Mertk* and *Tyro3*. **a**, Representative images of microcomputed tomography (μCT) of the metaphyseal proximal region of tibias from 8-week-old *Mertk^flox/flox^*, *Ctsk-cre^+^Mertk^flox/flox^* mice, *Tyro3^flox/flox^* and *LysM-cre^+^Tyro3^flox/flox^* female mice (longitudinal view of cancellous bone). **b,** Representative images and magnifications of TRAP/Hematoxylin staining of femur from *Mertk^flox/flox^*, *Ctsk-cre^+^Mertk^flox/flox^* mice, *Tyro3^flox/flox^* and *Ctsk-cre^+^Tyro3^flox/flox^* mice. **c,d,e,f,** Quantification of bone volume (BV/TV) (c), trabecular number (TbN) (d), trabecular thickness (TbTh) (e) and trabecular separation (TbSp) (f) of *Mertk^flox/flox^* and *Ctsk-cre^+^Mertk^flox/flox^* mice (n=13/9, mean±SEM, unpaired t-test). **g,h,** Histomorphometric analysis of osteoclast (N.Oc/B.PM) (g) and osteoblast number (N.Ob/B.PM) (h) of *Mertk^flox/flox^* and *Ctsk-cre^+^Mertk^flox/flox^* mice (n=6, mean±SEM, unpaired t-test). **i,j,k,l,** Quantification of bone volume (BV/TV) (i), trabecular number (TbN) (j), trabecular thickness (TbTh) (k) and trabecular separation (TbSp) (l) of *Tyro3^flox/flox^* and *Ctsk-cre^+^Tyro3^flox/flox^* mice (n=5/14, mean±SEM, unpaired t-test). **m,n,** Histomorphometric analysis of osteoclast (N.Oc/B.PM) (m) and osteoblast number (N.Ob/B.PM) (n) of *Tyro3^flox/flox^* and *Ctsk-cre^+^Tyro3^flox/flox^* mice (n=5/6, mean±SEM, unpaired t-test).

### Osteoclast precursor cell morphology regulated by MERTK dictates fusion capacity

In a next step we aimed to unravel the role of MERTK in osteoclast fusion and late osteoclast differentiation. We performed *ex vivo* osteoclast cultures of *Ctsk-cre^+^Mertk^flox/flox^* mice. Although we observed slightly decreased osteoclast numbers in *Ctsk-cre^+^Mertk^flox/flox^*mice, we observed highly increased osteoclast size and fusion number index indicating that loss of *Mertk* promotes osteoclast fusion (Fig. 5a-d). We could confirm these findings by pharmacological MERTK blockade using the small molecule inhibitor R992. Treatment of osteoclast cultures on day three (pre-fusion) led to increased fusion number index and osteoclast size (Suppl. Fig. 3a-c). Subsequently, we performed time-lapse microscopy of osteoclast fusion events. We used cells from *LysM-cre^+^Mertk^flox/flox^* mice to obtain cells with *Mertk*-deficiency also in early differentiation stages. We observed that osteoclast fusion is restricted in osteoclast precursor cells with spherical and round cell morphology with high contractility (Fig. 5e, red arrows) (Suppl. Video 6). We could not record any fusion event in cells displaying this morphology. Fusion-competent cells are characterized in contrast by a spread appearance with high numbers of filopodia and lamellipodia (Fig. 5e, green arrows and f) (Suppl. Video 7). Analysis of filopodia formation in *Mertk-*deficient osteoclast precursor cells by F-actin immunofluorescence staining confirmed increased filopodia formation (Fig. 5f and g).

**Figure 5.**
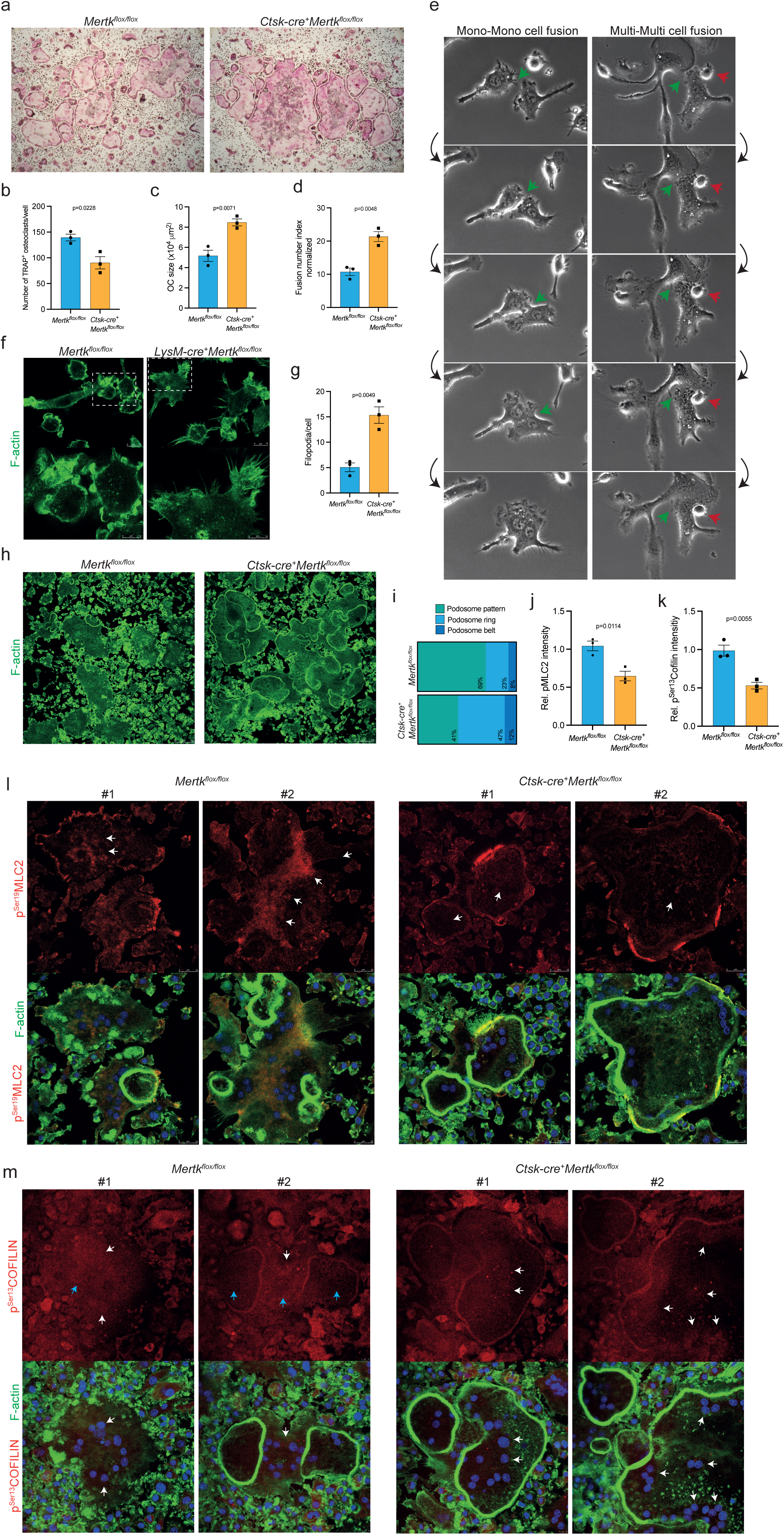
Deletion of Mertk enhances osteoclast fusion and promotes podosome ring formation. **a**, Representative images of *ex vivo* osteoclast differentiation assays from *Mertk^flox/flox^* and *Ctsk-cre^+^Mertk^flox/flox^* mice. **b,c,d,** Analysis of osteoclast number (b), osteoclast size (c) and fusion number index (d) in *ex vivo* osteoclast differentiation assays from *Mertk^flox/flox^* and *Ctsk-cre^+^Mertk^flox/flox^* mice (n=3, mean±-SEM, unpaired t-test). **e,** Time lapse microscopy images illustrating fusion events of mononuclear and multinuclear cells (green arrow displaying fusion events by fusion competent cells, red arrows displaying fusion-incompetent cells). **f,** Representative images of F-actin immunofluorescence stainings of filopodia formation in osteoclast precursor cells from *Mertk^flox/flox^* and *LysM-cre^+^Mertk^flox/flox^* mice. **g,** Quantification of filopodia per osteoclast precursor cell (n=3 random HPF, mean±SEM, unpaired t-test). **h,i,** Representative images (h) and analysis (i) of F-actin immunofluorescence stainings of podosome pattern in osteoclasts from *Mertk^flox/flox^* and *Ctsk-cre^+^Mertk^flox/flox^* mice. **j,k,** Analysis of immunofluorescence staining intensity of p^Ser19^MLC2 (J) and p^Ser3^COFILIN (K) in osteoclasts generated from *Mertk^flox/flox^* and *Ctsk-cre^+^Mertk^flox/flox^* mice (n=3 random HPF, mean±SEM, unpaired t-test). **l,m,** Representative images of F-actin and p^Ser19^MLC2 (l) / p^Ser3^COFILIN (m) immunofluorescence staining of osteoclast cultures from *Mertk^flox/flox^* and *Ctsk-cre^+^Mertk^flox/flox^* mice (#1 showing immature osteoclasts and #2 showing osteoclasts with higher maturity present in the indicated conditions).

### Central actomyosin contraction induced by MERTK-COFILIN-pMLC2 inhibits osteoclast maturation

After fusion, multinucleated osteoclasts undergo cytoskeletal reorganization for apico-basal polarization and sealing zone formation. Osteoclasts plated on glass display formation of podosome pattern which develop into podosome ring and subsequently into podosome belt upon maturation. We performed F-actin immunofluorescence staining of *Mertk-deficient* osteoclast cultures plated on glass. Analysis of podosome patterning in *Mertk-*deficient osteoclasts showed highly increased podosome ring formation, suggesting that loss of *Mertk* improves osteoclast maturation and function (Fig. 5h and i). Previous results showed that NMMIIA regulates osteoclast fusion, podosome organization and motility^35,36^. NMMIIA is controlled by phosphorylation of its myosin light chain^25^. We examined the localization of NMMIIA in osteoclasts under normal growth conditions. NMMIIA showed a strong and diffuse whole cell staining, colocalized with F-actin and slightly tended to be enriched at the cell border (Suppl. Fig. 4a). We examined spatial NMMIIA activation by staining for Ser19 phosphorylation of MLC2 in osteoclasts. p^Ser19^MLC2 did not co-localize with podosome clouds or podosome rings (Suppl. Fig. 4b, first panel). In contrast, we observed high p^Ser19^MLC2 activity co-localizing with osteoclast adhesion structures (Suppl. Fig. 4b, second panel, arrows). Interestingly, we found osteoclasts with podosome belts not interconnected with adhesion structures. In this case they lacked p^Ser19^MLC2 activity (Suppl. Fig. 4b). When podosome belts were integrated into adhesion structures they showed strong p^Ser19^MLC2 activity (Suppl. Fig. 4b, third panel, arrows). Therefore, we hypothesized that spatial localization and activation of actomyosin forces in the cell periphery of osteoclasts adjacent to cell-matrix contacts are key for the formation and maintenance of the osteoclast sealing zone. We performed immunofluorescence staining of p^Ser19^MLC2 in *Mertk*-deficient osteoclasts. Immature control osteoclasts defined by displaying podosome pattern exhibited high central myosin light chain phosphorylation, which was reduced by deletion of *Mertk* (Fig. 5j and l (#1), arrows). Likewise, osteoclasts with higher maturity defined by podosome belt formation induced by *Mertk* display decreased p^Ser19^MLC2, indicating that reduced central actomyosin contraction ameliorates osteoclast podosome patterning leading to actin ring formation (Fig. 5l (#2), arrows). It was shown that active COFILIN promotes podosome belt formation in osteoclasts^37^. Therefore, we stained for COFILIN inactivating Ser3 phosphorylation and observed that increased osteoclast maturation induced by deletion of *Mertk* correlated with decreased p^Ser3^COFILIN staining intensity (Fig. 5k, m). Immature control osteoclasts showed high levels of p^Ser3^COFILIN, whereas cellular regions inside podosome rings exhibited low levels of p^Ser3^COFILIN (Fig. 5m, blue arrows). Especially *Mertk*-deficient osteoclasts with podosome belt formation showed low p^Ser3^COFILIN signal (Fig. 5m (#2). Interestingly, we observed that p^Ser3^COFILIN was correlated to the perinuclear region of osteoclasts (Fig. 5m, white arrows). Further observations by immunofluorescence and live-cell imaging revealed a shift of the nuclei to the osteoclast cell border forming circular syncytia upon maturation and this developmental process was enhanced by deletion of *Mertk* (Suppl. Video 8) and inhibited by PROS1 (Suppl. Video 9), indicating that spatial MERTK-COFILIN activation prevents intracellular nuclear migration and pulling of the nucleus to the cell borders (Fig. 5k and m, arrows) (Suppl. Fig. 2g, arrows). The correlation of high spatial p^Ser3^COFILIN with the perinuclear region is in line with high MERTK expression in the nucleus (Fig. 2c). Interestingly, although upstream regulator of both MLC2 and COFILIN is in principle RHOA, it seems that activation of these downstream molecules is uncoupled in osteoclasts, as MLC2 activation does not correlate with presence of nuclei. In line with that, we observed that MERTK mainly phosphorylates COFILIN in osteoclasts, whereas its activating effects on RHOA were masked by TYRO3-mediated silencing of RHOA.

### Peripheral actomyosin contraction inhibited by TYRO3 induces osteoclast maturation

Next, we performed *ex vivo* osteoclast cultures of *Ctsk-cre^+^Tyro3^flox/flox^*mice. Osteoclast numbers were increased upon deletion of *Tyro3* (Fig. 6a and b). F-actin immunofluorescence staining showed increased podosome belt formation in osteoclasts with deletion of *Tyro3* in contrast to *Mertk*-deficiency osteoclasts which induced increased podosome ring formation (Fig. 6c and d). We observed likewise to the previous results in osteoclast precursors (Figure 3a) that increased RHOA activation in mature osteoclasts was induced by deletion of *Tyro3* and PROS1 could not decrease RHOA activation in absence of TYRO3, indicating that TYRO3 inhibits RHOA activity in osteoclasts (Fig. 6e). Concomitantly, overall p^Ser19^MLC2 staining intensity was increased in *Tyro3*-deficient osteoclasts (Fig. 6f). Further morphologic analysis revealed that control osteoclasts were mostly spread and round to rectangular with relatively smooth and thin actin rich peripheral cell boarders. *Tyro3*-deficient osteoclasts displayed a retracted, voluminous and mostly polarized appearance (Fig. 6h) (Suppl. Video 10). Most prominently *Tyro3* deficiency led to membrane protrusions with multiple foot process like formations at the cell periphery, suggesting improved cell-matrix interaction (Fig. 6h). *Tyro3* knockout osteoclasts exhibited high p^Ser19^MLC2 signals in the cell periphery, located in high proximity to adhesion structures. We analyzed the fluorescence intensity of p^Ser19^MLC2 in peripheral areas (defined as 10% of diameter from cell edge) relative to the inner 90% area (central signal). The ratio of peripheral signal in relation to central signal was highly increased in osteoclasts with deletion of *Tyro3* in comparison to controls (Fig. 6g). Our data suggest that loss of *Tyro3* contributes to an activated osteoclast phenotype with increased actin ring formation induced by activation of RHOA-mediated spatial actomyosin contraction, especially in peripheral cell compartments, adjacent to osteoclasts adhesion structures. For bone resorption, osteoclasts initiate a sealing zone, reduce their adhesive surface, and consequently round up a process called apicobasal polarization^23^. The voluminous appearance of *Tyro3-*deficient osteoclasts led us hypothesize that increased RHOA activation levels induced by loss of *Tyro3* promotes apicobasal polarization of osteoclasts. The role of RHOA in regulating osteoclast cytoskeletal configuration is controversial^38^. Our model suggests that spatially well-ordered RHOA activation at adhesion structures improves osteoclast biomechanotransduction to induce sealing zone formation and enhance osteoclast function. In turn, MERTK-RHOA-COFILIN mediated central actomyosin contraction is detrimental for podosome patterning and subsequently osteoclast activation.

**Figure 6:**
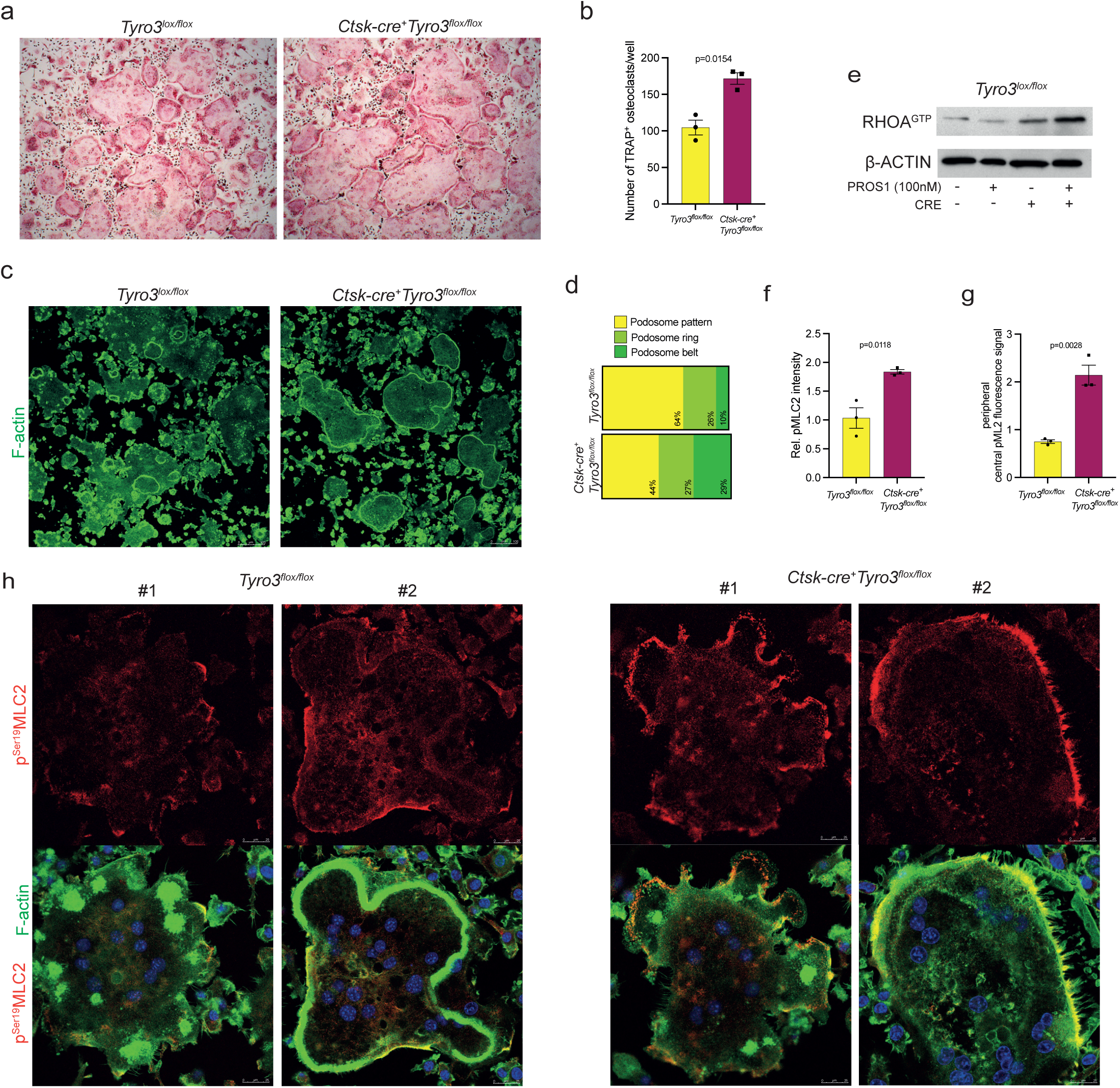
*Tyro3* controls spatial phosphorylation of Non-Muscle Myosin II regulatory light chain which regulates podosome belt formation in osteoclasts. **a**, Representative images of *ex vivo* osteoclast differentiation assays from *Tyro3^flox/flox^* and *Ctsk-cre^+^Tyro3^flox/flox^* mice. **b,** Analysis of osteoclast number in *ex vivo* osteoclast differentiation assays from *Tyro3^flox/flox^* and *Ctsk-cre^+^Tyro3^flox/flox^* mice (n=3, mean±SEM, unpaired t-test). **c,d,** Representative images (c) and analysis (d) of podosome pattern by F-actin immunofluorescence staining in osteoclasts from *Tyro3^flox/flox^* and *Ctsk-cre^+^Tyro3^flox/flox^* mice. **e,** Immunoblot analysis of RHOA signaling in osteoclasts harboring *Tyro3* knockout and treated with PROS1. **f,g,** Analysis of total (f) and peripheral/central (g) p^Ser19^MLC2 staining intensity in osteoclasts from *Tyro3^flox/flox^* and *Ctsk-cre^+^Tyro3^flox/flox^* mice (n=3 random HPF, mean±SEM, unpaired t-test). **h,** Representative images of F-actin and p^Ser19^MLC2 immunofluorescence stainings of immature (#1) and mature (#2) osteoclasts from *Tyro3^flox/flox^* and *Ctsk-cre^+^Mertk^flox/flox^* mice.

### Diminution and aggravation of osteolytic bone disease of breast cancer bone metastases by lysozyme M-mediated deletion of *Mertk* and *Tyro3*

To investigate a potential role of MERTK in cancer cell–osteoclast interaction in order to elucidate if MERTK represents a novel target to inhibit osteolysis in breast cancer bone metastasis, we used the syngeneic EO771 breast cancer cell line and injected luciferase transduced EO771 cells into *Mertk^flox/flox^*and *LysM-cre^+^Mertk^flox/flox^* C57BL/6J mice. Metastatic spread was monitored by bioluminescence imaging. Bone metastasis progression in hind limbs was not significantly changed after 12 days (Fig. 7a and b). Analysis of metaphyseal cancellous bone by μCT showed increased bone volume, trabecular number and trabecular thickness in EO771 tumor-bearing *LysM-cre^+^Mertk^flox/flox^* mice in comparison to *Mertk^flox/flox^* mice, suggesting that MERTK mediates osteolysis in breast cancer bone metastases (Fig. 7c-h). We performed TRAP/Hematoxylin staining of tibia and observed decreased osteoclast number on the surface of trabecular bone of tumor-bearing *LysM-cre^+^Mertk^flox/flox^* mice, whereas osteoblast numbers were not significantly changed (Fig. 7i-k). Furthermore, we observed decreased intratumoral TRAP^+^ mononuclear cells, indicating decreased intratumoral osteoclast precursor cell recruitment induced by deletion of *Mertk* (Fig. 7l). Next, we injected EO771 cells into *Tyro3^flox/flox^* and *LysM-cre^+^Tyro3^flox/flox^*C57BL/6J mice. Again, we did not observe differences in hind limb tumor load in comparison to EO771 bearing *Tyro3^flox/flox^* control mice (Fig. 7m and n). In turn, μCT analysis showed decreased bone volume and trabecular number in EO771 tumor-bearing *LysM-cre^+^Tyro3^flox/flox^*mice in comparison to *Tyro3^flox/flox^* mice after 12 days (Fig. 7o-t). We observed increased osteoclast numbers on the bone surface, whereas osteoblast numbers were not changed (Fig. 7u-w). This was corroborated by increased intratumoral TRAP^+^ mononuclear cells, indicating increased osteoclast precursor cell recruitment induced by deletion of *Tyro3* (Fig. 7x).

**Figure 7:**
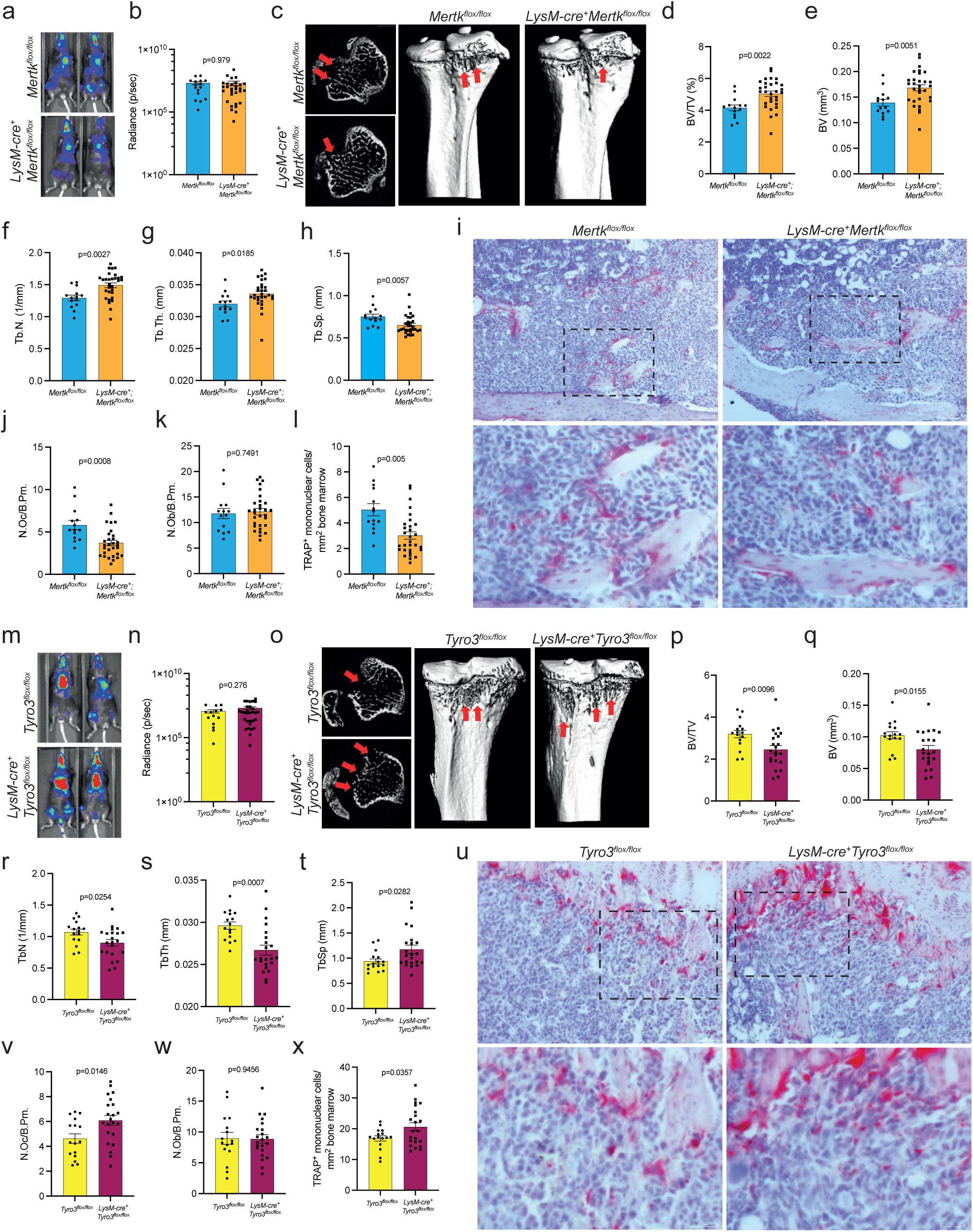
Diminution and aggravation of osteolytic bone disease of breast cancer bone metastases by *lysozyme M* - mediated deletion of *Mertk* and *Tyro3.* **a**, Luciferase^+^ EO771 breast cancer cells were injected intracardially in *Mertk^flox/flox^* and *LysM-cre^+^Mertk^flox/flox^* mice. **b,** Analysis of tumor load in tibias by Bioluminescence Imaging after 10 days (n=16/30, mean±SEM, unpaired t-test). **c,d,e,f,g,h,** Representative μCT images and 3D reconstructions (c) and analysis of BV (d), BV/TV (e), Tb.N (f), Tb.Th (g) and Tb.Sp (h) of trabecular bone of metaphyseal proximal region of the tibia of *Mertk^flox/flox^* and *LysM-cre^+^Mertk^flox/flox^* mice (n=16/30, mean±SEM, unpaired t-test). **i,j,k,l,** Representative images (i) and analysis of osteoclast number (N.Oc/B.Pm) (j), osteoblast number (N.Ob/B.Pm) (k) and osteoclast precursor cell number (TRAP^+^ mononuclear cells/mm^2^ bone marrow) (l) of TRAP/hematoxylin staining of tibia from *Mertk^flox/flox^* and *LysM-cre^+^Mert-k^flox/flox^* mice injected with EO771 cells. **m,** Luciferase^+^ EO771 breast cancer cells were injected intracardially in *Tyro3^flox/flox^* and *LysM-cre^+^Ty-ro3^flox/flox^* mice. **n,** Analysis of tumor load in tibias by Bioluminescence Imaging after 10 days (n=14/30, mean±SEM, unpaired t-test). **o,p,q,r,s,t,** Representative μCT images and 3D reconstructions (o) and analysis of BV (p), BV/TV (q), Tb.N (r), Tb.Th (s) and Tb.Sp (t) of trabecular bone of metaphyseal proximal region of the tibia of *Tyro3^flox/flox^* and *LysM-cre^+^Tyro3^flox/flox^* mice (n=16/22, mean±SEM, unpaired t-test). **u,v,w,x,** Representative images (u) and analysis of osteoclast number (v), osteoblast number (w) and osteoclast precursor cell number (x) of TRAP/hematoxylin staining of tibia from *Tyro3^flox/flox^* and *LysM-cre^+^Tyro3^flox/flox^* mice injected with EO771 cells (n=16/22, mean±-SEM, unpaired t-test).

As expected, injection of EO771 breast cancer cells into mice harboring *Ctsk-cre* mediated deletion of *Mertk* or *Tyro3* did not affect tumor load but led to decreased bone volume and increased osteolytic bone destruction in line with the inhibitory function of MERTK as well as TYRO3 in osteoclasts (Suppl. Fig. 5a-p).

Together these findings indicate that TAM receptor signaling represents a novel biological regulatory system in the tumor microenvironment of breast cancer bone metastases controlling osteolytic bone destruction with MERTK promoting cancer-induced osteoclast formation, whereas TYRO3 exerts bone protective effects.

## Discussion

In this study, we could demonstrate that MERTK and TYRO3 represent potent regulators of osteoclastic bone remodeling. The role of MERTK in osteoclasts is complex and context-dependent. Mice with *lysozyme M* mediated myeloid cell-specific knockout of *Mertk* exhibited high bone mass, whereas *cathepsin K* mediated osteoclast-specific deletion of *Mertk* resulted in an osteoporotic bone phenotype. MERTK represents an osteoclast-intrinsic regulatory factor enhancing osteoclast formation but inhibiting osteoclast fusion and late osteoclast maturation and function. Mechanistically, MERTK is critical to maintain amoeboid migration mode in osteoclast precursors mediating osteoclast motility leading to increased osteoclast formation. We found that RANKL downregulates *Mertk* expression in osteoclast precursors. RANKL is a cell membrane-bound protein. It can be cleaved into its soluble form by proteases, but soluble RANKL does not contribute to normal bone homeostasis^39^. Osteocytes are the main source of RANKL and can reach the bone surface with their dendritic processes^40,41^. We hypothesize, after MERTK-mediated osteoclast precursor cell migration to targeted bone resorption sites, its expression is downregulated by osteocytes, the main source of RANKL, at the bone surface. Thereby, osteoclast fusion and cytoskeletal reorganization to induce sealing zone formation is initiated. We could confirm these hypotheses by using *cathepsin K* mediated osteoclast-specific deletion of *Mertk,* which resulted in an osteoporotic bone phenotype. Our results suggest that MERTK induces mainly central actomyosin contractility in osteoclast precursors via RHOA-COFILIN pathway, thereby inhibiting actin polymerization to maintain round, spherical cell shape to promote amoeboid migration mode with high velocity. In mature osteoclasts MERTK signaling inhibited osteoclast podosome ring formation. Other authors showed that RHOA enhances osteoclastogenesis via mTOR-Nfatc1 and conditional knockout of RHOA resulted in increased bone mass^42,43^. In contrast, another study showed that activation of RHOA led to podosome belt dissolution and microtubule deacetylation thereby inhibiting osteoclast maturation^38^. Furthermore, it was shown that COFILIN is connected to podosome belt assembly. Active COFILIN is localized in podosome core structures, whereas phosphorylated, inactive cofilin is concentrated in the immature podosome clouds^37^. Our results suggest that MERTK could promote RHOA mediated COFILIN inactivation in osteoclasts, thereby inhibiting podosome ring formation. Additionally, MERTK may mediates RHOA - mammalian Diaphanous (mDIA)-related formin - induced microtubule deacetylation to inhibit osteoclast maturation. The osteoclast promoting function of RHOA in early phases via mTOR-Nfatc1 and inhibiting function of RHOA-COFILIN/mDIA in late osteoclast maturation is in line with our obtained results.

RANKL also downregulates TYRO3 which in comparison to MERTK likewise promotes an osteoclast conducive cytoskeletal configuration to induce sealing zone formation, although TYRO3 inhibits RHOA-ROCK signaling. We observed that in mature osteoclasts, deletion of *Tyro3* promoted late osteoclast differentiation and maturation by coupling podosome belts with osteoclast adhesion structures at the cell periphery via RHOA-MLC2 actomyosin mediated improved biomechanotransduction. We speculate that TYRO3 controls an unknown plasma membrane located RHO-GAP, which silences RHOA locally at the plasma membrane, whereas MERTK activates cytoplasmic RHOA leading to COFILIN phosphorylation. We suggest that RHOA activated at the plasma membrane fosters osteoclast sealing zone formation, whereas cytoplasmic RHOA inhibits these processes, and these spatial processes are controlled by MERTK and TYRO3. In line with these hypotheses, we observed that extracellular TAM receptor ligand PROS1 inhibited RHOA via TYRO3, a process taking place obligately at the cell membrane. MERTK might exhibit an unknown intracellular function leading to cytoplasmic RHOA-COFILIN signaling. In line with that we observed high MERTK signal in the nucleus in osteoclasts.

The osteoporotic bone phenotype induced by osteoclast-specific deletion of *Mertk* as well as *Tyro3* revealed a thus far unknown negative regulatory role of TAM receptors in osteoclasts. In concordance, case reports showed that children with inherited PROS1 deficiency exhibited osteopenia which led to development of pathological fractures^44,45^. In addition, it was recently shown that osteoporosis is connected to a single-nucleotide polymorphism (SNP) associated with the PROS1 gene, raising the intriguing possibility that PROS1 and TAM receptors might represent a clinical-relevant regulatory system in bone homeostasis contributing to the pathogenesis of bone diseases such as osteoporosis via inhibition of osteoclasts^46^.

Our results contradict several previous studies suggesting an osteoclast promoting function of TYRO3^11–15^. Mice with *Tyro3* germline deletion exhibited high bone mass, reduced osteoclast numbers and reduced RA-induced bone erosions^13^. These data suggest that similar to MERTK, TYRO3 induces early osteoclast differentiation, but inhibits late osteoclast maturation and bone resorption. Another reason could be that *Tyro3* germline deletion might influence other cell types in the bone marrow microenvironment leading to expression of osteoclast-promoting factors. In line with our results, one study showed that soluble TYRO3 correlates with RA disease activity score and treatment of osteoclast cultures with soluble TYRO3 led to increased osteoclast differentiation^16^. Knockout of *Tyro3* increased osteoclast precursor cell motility. We obtained similar results in migration of osteoblasts^9^. Nevertheless, interestingly activation of TYRO3 promoted mesenchymal-like migration, a migration mode leukocytes are dependent on when migrating in 2D environments like blood vessels, in contrast to 3D migration in the interstitium where leukocytes are dependent on amoeboid migration^47^. Therefore, our results do not exclude that TYRO3 might be a pro-osteoclastogenic factor in circulating monocytes for diapedesis and blood vessel extravasation, which is dependent on cytoskeletal reorganization and is crucial in autoinflammatory diseases such as RA^13,15,48,49^

In contrast to RA, breast cancer metastases are characterized by an anti-inflammatory tumor microenvironment^50,51^. In the mouse model of breast cancer bone metastasis, we observed that conditional deletion of *Mertk* reduces breast cancer associated osteolytic bone disease. We suggest that in our model the tumor associated anti-inflammatory setting induces MERTK-mediated amoeboid migration of bone marrow macrophages towards tumor adjacent bone resorption sites to induce osteolytic bone disease. In contrast, deletion of *Tyro3* in myeloid cells increased osteoclast precursor cell migration and induced osteolytic bone disease by increasing osteoclast formation and osteoclastic bone resorption. Injection of breast cancer cells into mice harboring osteoclast-specific deletion of *Mertk* and *Tyro3* resulted in aggravated osteolytic bone diseases confirming the inhibiting function of MERTK as well as TYRO3 in osteoclast maturation and function.

In summary, in the context of cancer-induced osteolytic bone disease activation of MERTK suppresses and transforms osteoblasts, counteracting bone formation and recruiting osteoclast precursors to generate osteoclasts and mediate osteolysis. Hence, MERTK is a target in the tumor microenvironment to inhibit osteolytic bone metastasis.

As our discoveries do not seem to be tumor-specific, the discovery of the function of MERTK in the bone could have deep implications for the treatment of various bone diseases such as osteoporosis.

## Materials & Methods

### Osteoclast cultures

Femur and tibia of 8-14-week-old mice were flushed with PBS and bone marrow cells were seeded out in on 5 cm Petri dishes with standard α-MEM Medium (α-MEM GlutaMAX) supplemented with 10% FBS and 1% Penicillin/Streptomycin. After 3 h suspension cells were transferred to 10 cm Petri dish and cultured for 72 h with 100ng/ml M-CSF to generate bone marrow macrophages. Bone marrow macrophages were trypsinized and transferred to 96-well cell culture plates (10.000 cells/well) and treated with 30 ng/ml M-CSF and different concentrations of RANKL (10 ng/ml or 50 ng/ml). Medium was replaced after 72 h. Multinucleated cells were observed after another 96 h – 120 h.

### Cell migration analysis

Cell tracking was performed manually using the ImageJ Manual Tracking plugin developed by Cordelieres. After tracking the Chemotaxis and Migration Tool (Ibidi, GmbH, https://ibidi.com/chemotaxis-analysis/171-chemotaxis-and-migration-tool.html) was used to calculate migration parameters.

### Murine breast cancer cell line cell culture

EO771 (the gift from Massimiliano Mazzone) cells were cultured in DMEM medium supplemented with 10% fetal bovine serum (FBS) and 1% penicillin/streptomycin (P/S).

### RNA isolation/RT-qPCR

RNA was isolated using Fisher Scientific Invitrogen PureLink RNA Kit. cDNA synthesis was performed using a Thermo Scientific First strand cDNA synthesis kit. Quantitative RT-PCR was performed using Taqman GeneExpression Assays (ThermoFisher) and Eppendorf MasterCycler technology. Copy numbers were calculated using the standard curve method for absolute quantification.

### Immunoblot

For harvest, cells were washed in ice-cold PBS and lysed in RIPA Buffer (Thermo Fisher), containing PhosSTOP phosphatase inhibitor cocktail (Roche) and protease inhibitor cocktail (Roche). After SDS-PAGE proteins were electrotransferred to nitrocellulose membranes and blocked in 5% bovine serum albumin for 1 h. Blocked membranes were probed with primary antibodies overnight at 4 °C in the same buffer, followed by secondary antibody conjugated to HRP in blocking solution for 1 h shaking at room temperature. Bands were visualized using enhanced chemiluminescence (ECL) detection. GTP-bound RHOA was analyzed by performing a pull-down assay with Cell Biolabs, Inc. Rho Activation Assay kit.

### Immunofluorescence imaging

Bone marrow macrophages were cultured with 30 ng/ml M-CSF and 50 ng/ml RANKL on glass coverslips for 48 h to generate osteoclast progenitor cells. Bone marrow macrophages were cultured with 30 ng/ml M-CSF and 50 ng/ml RANKL on glass coverslips for 5-7 days to generate multinucleated osteoclasts.

Cells were fixed for 10 min in 4% PFA and permeabilized for 3-5 min with 0.01% TRITON X-100. Blocking was performed for 30 min with 1% BSA in PBS with Tween (PBS-T).

F-actin was visualized by incubating the cells with Alexa Fluor 488 Phalloidin (Invitrogen) (1:50) in a blocking solution for 30 min at room temperature in the dark.

Anti-pMLC2 (Ser19) antibody #3671 (Cell Signaling) (1:100), anti-pCOFILIN antibody #3313 (Cell Signaling) (1:100) or anti-NMMllA antibody #3403 (Cell Signaling) (1:100) was incubated as the first antibody (1:100) for 1 h at room temperature in blocking solution. After rigorous washing, secondary antibody goat anti-rabbit AlexaFluor 555 (Thermo Fisher) (1:200) was incubated together with Alexa Fluor 488 Phalloidin (Invitrogen) (1:50) and DAPI (1:500) in blocking solution for 1 h at room temperature in the dark. After rigorous washing, cells were mounted in Vectashield mounting medium (Vector Laboratories).

Confocal microscopy was conducted using the Leica TCS SP8 X microscope (Software Leica LAS X). Immunofluorescence staining intensity was measured using ImageJ.

### Generation of luciferase expressing breast cancer cell line

EO771-Luc were generated by lentivirally transducing them with the firefly luciferase gene. The third generation HIV-1 derived lentiviral vector LeGO-iG2-Puro^+^-Luc2 expresses the Luc2 variant of firefly luciferase (cloned from AddGene Plasmid 24337) under the control of an SFFV-promoter, linked by an IRES to a second open reading frame consisting of eGFP, a 2A-peptide and the puromycin resistance PAC^53^. After lentiviral transduction, luciferase^+^ cells were selected by Puromycin treatment (1 μg/ml) and purity >99% was confirmed by FACS analysis. Sufficient luciferase signal of the cells was validated in vitro by several dilution series, luciferin treatment, and image acquisition with IVIS 200 imaging system (PerkinElmer).

### μCT

μCT was used for 3D analyses of long bones. Long bones of mice were analyzed using high-resolution μCT with a fixed isotropic voxel size of 10 μm (70 peak kV at X μA 400 ms integration time; Viva80 microCT; Scanco Medical). All analyses were performed on digitally extracted bone tissue using 3D distance techniques (Scanco Medical), as reported previously. Region of interest (ROI) was defined manually by drawing contours in slices.

### Histology

Bones were fixed with 4% paraformaldehyde for 48 h and the tibia was decalcified with EDTA. Paraffin blocks were cut into 5-μm-thick sections. Two non-serial sections of each bone were assessed. TRAP staining was performed for 30 min followed by nuclear counterstaining with hematoxylin. The number of osteoclasts and osteoblasts at the bone surface was measured using the osteomeasure system (Osteometrics).

### Statistics and reproducibility

Data were means ± SEMs. Statistical significance was determined by a two-tailed unpaired t-test, unless otherwise stated. Significant outliers were calculated and excluded from the analysis. All statistical analyses were performed using GraphPad Prism 5 software. For data presentation, a representative experiment was chosen and included in the manuscript.

### Key resources table

| REAGENT or RESOURCE | SOURCE | IDENTIFIER |
| --- | --- | --- |
| **Antibodies** |  |  |
| Rabbit monoclonal anti-Cofilin | Cell Signaling | Cat#D3F9 |
| Rabbit monoclonal anti-phospho-Cofilin (Ser3) | Cell Signaling | Cat#77G2 |
| Phospho-Myosin Light Chain 2 (Ser19) | Cell Signaling | Cat#3671 |
| Alexa Fluor 488 Phalloidin | Thermo Fisher | Cat#A12379 |
| Myosin IIa | Cell Signaling | Cat#3403 |
| Goat anti-Rabbit IgG (H+L) Cross-Adsorbed Secondary Antibody, Alexa Fluor 555 | Thermo Fisher | Cat #A-21428 |
| **Biological samples** |  |  |
| Mouse: bone marrow harvested from C57BL/6J & indicated conditional knockout mice | This paper | N/A |
| Mouse: Tibia and femur harvested from indicated conditional knockout mice | This paper | N/A |
| **Chemicals, peptides, and recombinant proteins** |  |  |
| Human plasma purified Protein S | Enzyme Research Laboratories | Cat#HPS |
| R992<sup>9</sup> | RIGEL inc. | N/A |
| Human recombinant M-CSF | Peptotech | Cat#310-01 |
| Human recombinant soluble RANKL | Peptotech | Cat#300-25 |
| **Critical commercial assays** |  |  |
| RhoA Activation Assay Kit | Cell Biolabs, INC | Cat#STA-403-A |
| Taqman Universal PCR-Mastermix | Thermo Fisher | Cat#4304437 |
| RevertAid First Strand cDNA synthesis Kit | Thermo Fisher | Cat#K1621 |
| BlackPREP Rodent Tail DNA Kit | IST Innuscreen | N/A |
| PureLink RNA Mini Kit | Thermo Fisher | Cat#12183020 |
| **Experimental models: Cell lines** |  |  |
| EO771 | M. Mazzone | N/A |
| EO771-luc2 | Our lab | Engelmann et.al.<sup>9</sup> |
| **Experimental models: Organisms/strains** |  |  |
| Mouse: C57BL/6J | FTH UKE |  |
| Mouse: Mertk<sup>flox/flox</sup> | C. Rothlin | Fourgeaud et. al.<sup>8</sup> |
| Mouse: Tyro3<sup>flox/flox</sup> | C. Rothlin | Engelmann et.al.<sup>9</sup> |
| Mouse: LysM-cre | H. Tapaleenmäki | Strain #:004781 |
| Mouse: Ctsk-cre | H. Tapaleenmäki | Nakamura et. al.<sup>52</sup> |
| **Oligonucleotides** |  |  |
| Mertk | Thermo Fisher | Mm07295861_m1 |
| Tyro3 | Thermo Fisher | Mm00444557_g1 |
| Pros1 | Thermo Fisher | Mm01343426_m1 |
| Gas6 | Thermo Fisher | Mm00490378_m1 |
| Nfatc1 | Thermo Fisher | Mm01265944_m1 |
| DcStamp | Thermo Fisher | Mm04209236_m1 |
| Acp5 | Thermo Fisher | Mm00475698_m1 |
| Ctsk | Thermo Fisher | Mm00484039_m1 |
| Recombinant DNA |  |  |
| Lentiviral vector LeGO-iG2-Puro<sup>+</sup>-Luc2 | Kristoffer Riecken | https://www.lentigo-vectors.de |
| Luc2 variant of firefly luciferase | AddGene | Plasmid 24337 |
| Software and algorithms |  |  |
| ImageJ | Publicly available | https://imagej.nih.gov/ij/ |
| Prism v8.0 | GraphPad Software | https://www.graphpad.com |
| microCT analysis software | Scanco Medical | N/A |
| Osteomeasure system | Osteometrics | N/A |
| Other |  |  |
| Bio-Rad CFX96 Real Time System | Bio-Rad | N/A |
| Eppendorf Master Cycler | Eppendorf | N/A |
| Viva80 microCT | Scanco Medical | N/A |

## Acknowledgments

This work was supported by the priority program μbone from the DFG (LO1863/5-1 to J.E., LO1863/5-1 to S.L., BE6658/1-1 and BE6658/2-1 to I.B.B.). S.L. was supported by a Heisenberg professorship (DFG) and is currently supported by the European Research Council (ERC) under the European Union’s Horizon 2020 research and innovation program (Grant Agreement No. 758713) and by the Hector Stiftung II. E.H. and H.T. received funding from the German Research Foundation (SPP 2084 to E.H. (HE 5208/5-1) and H.T (TA 1154/2-1) and Emmy Noether Program to H.T. (TA 1154/1-1 and TA 1154/1-2)). The authors are deeply grateful to Michael Horn from the UCCH *in vivo* optical imaging core facility for help with the intracardial injections and imaging of our mouse models. Confocal microscopy was conducted at the UKE Microscopy Imaging Facility (DFG Research Infrastructure Portal: RI_00489). We thank Bao-Uyen Huynh on behalf of the Animal Facility (FTH) at University Hospital Hamburg-Eppendorf for taking care of the mice housing and breeding.

## Author Contributions

Conceptualization, J.E., I.B.B. and S.L.; Methodology, J.E., I.B.B. and S.L.; Investigation, J.E., J.Z., M.S., C.M., M.H., D.R., K.R.; Writing – Original Draft, J.E.; Writing – Review & Editing, S.L. and I.B.B.; Funding Acquisition, I.B.B. and S.L.; Resources, E.A., S.G., C.R., K.P., C.B., E.H., H.T., S.L.; Supervision, I.B.B. and S.L.

## Declaration of interests

The authors declare no competing interests.

## Notes

### Competing Interest Statement

The authors have declared no competing interest.

