## Supplementary material for "TAM receptors control actomyosin dynamics in osteoclasts via RHOA-COFILIN-MYOSIN II signaling": Suppl. Figures

### Supplemental Figure 1

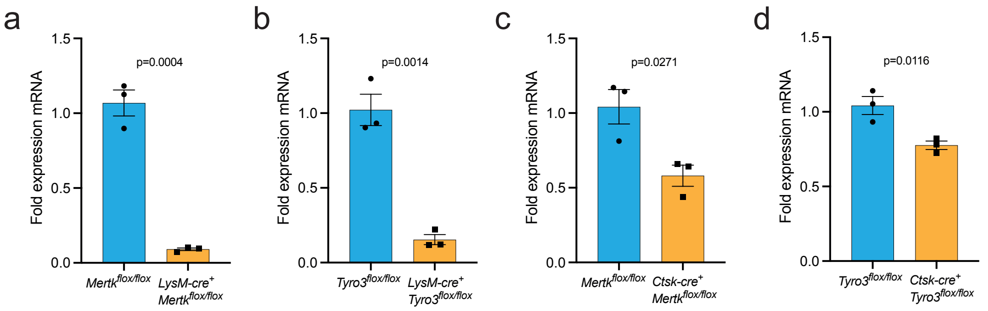


**Supplemental Figure 1. LysM-cre and Ctsk-cre mediated deletion of Mertk and Tyro3. a,** Analysis of *Mertk* expression in *ex vivo* osteoclast cultures from *LysM-cre*$^{+}$*Mertk*$^{flox/flox}$ mice (n=3, mean±SEM, unpaired t-test). **b,** Analysis of *Tyro3* expression in *ex vivo* osteoclast cultures from *LysM-cre*$^{+}$*Tyro3*$^{flox/flox}$ mice (n=3, mean±SEM, unpaired t-test). **c,** Analysis of *Mertk* expression in *ex vivo* osteoclast cultures from *Ctsk-cre*$^{+}$*Mertk*$^{flox/flox}$ mice (n=3, mean±SEM, unpaired t-test). **d,** Analysis of *Tyro3* expression in *ex vivo* osteoclast cultures from *Ctsk-cre*$^{+}$*Tyro3*$^{flox/flox}$ mice (n=3, mean±SEM, unpaired t-test).

### Supplemental Figure 2

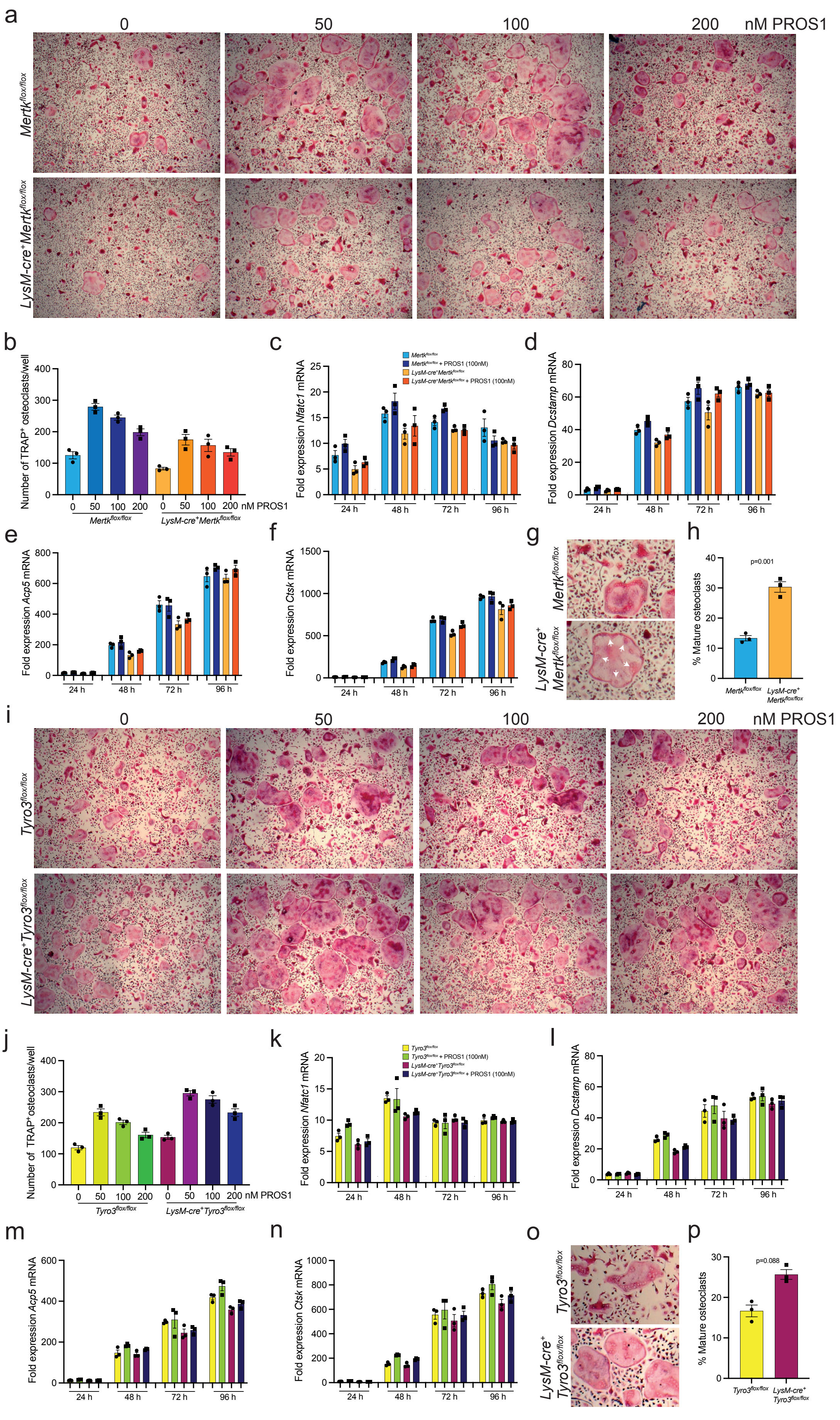


**Supplemental Figure 2. Effect of *lysozyme M* - mediated deletion of *Mertk* and *Tyro3* on osteoclast differentiation *in vitro*. a,b,** Representative images (a) and osteoclast numbers (TRAP$^{+}$ multinuclear cells with >3 nuclei) (b) of *ex vivo* osteoclast differentiation assays from *Mertk$^{flox/flox}$* and *LysM-cre$^{+}$Mertk$^{flox/flox}$* mice treated with different doses of PROS1 (50-200 nM), 30 ng/ml M-CSF and 50 ng/ml RANKL (n=3, mean±SEM). **c,d,e,f,** Evaluation of osteoclast differentiation marker *Nfatc1* (c), *Dcstamp* (d), *Acp5* (e) and *Ctsk* (f) by RT-qPCR in *Mertk$^{flox/flox}$* and *LysM-cre$^{+}$Mertk$^{flox/flox}$* osteoclast cultures treated with 100 nM PROS1, 30 ng/ml M-CSF and 50 ng/ml RANKL (n=3, mean±SEM). **g,** Magnifications showing osteoclast morphology in osteoclast cultures from *Mertk$^{flox/flox}$* and *LysM-cre$^{+}$Mertk$^{flox/flox}$* mice. White arrows depicting shift of nuclei to cell boundaries. **h,** Quantification of percentage of mature osteoclasts exhibiting typical osteoclast morphology in osteoclast cultures from *Mertk$^{flox/flox}$* and *LysM-cre$^{+}$Mertk$^{flox/flox}$* mice (n=3, mean±SEM). **i,j,** Representative images (i) and osteoclast numbers (TRAP$^{+}$ multinuclear cells with >3 nuclei) (j) of *ex vivo* osteoclast differentiation assays from *Tyro3$^{flox/flox}$* and *LysM-cre$^{+}$Tyro3$^{flox/flox}$* mice treated with different doses of PROS1 (50-200 nM), 30 ng/ml M-CSF and 50 ng/ml RANKL. **k,l,m,n** Evaluation of osteoclast differentiation marker *Nfatc1* (k), *Dcstamp* (l), *Acp5* (m) and *Ctsk* (n) by RT-qPCR in *Tyro3$^{flox/flox}$* and *LysM-cre$^{+}$Tyro3$^{flox/flox}$* osteoclast cultures treated with 100 nM PROS1 and 50 ng/ml RANKL (n=3, mean±SEM). **o,** Magnifications showing osteoclast morphology in osteoclast cultures from *Tyro3$^{flox/flox}$* and *LysM-cre$^{+}$Tyro3$^{flox/flox}$* mice. **p,** Quantification of percentage of mature osteoclasts exhibiting typical osteoclast morphology in osteoclast cultures from *Tyro3$^{flox/flox}$* and *LysM-cre$^{+}$Tyro3$^{flox/flox}$* mice (n=3, mean±SEM).

### Supplemental Figure 3

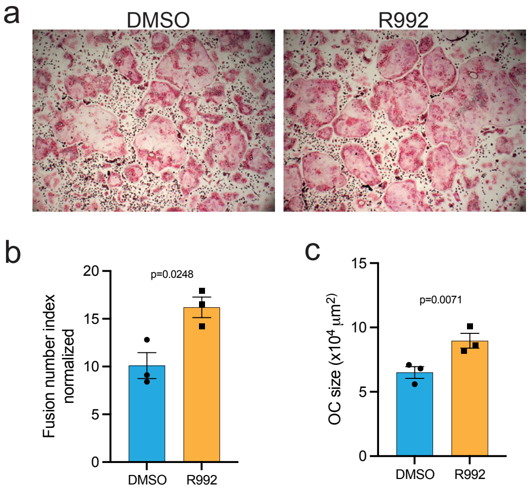


**Supplemental Figure 3. Pharmacologic MERTK blockade induces osteoclast fusion. a,b,c,** Representative images (a) and analysis of osteoclast fusion number index (b) and osteoclast size (c) of *ex vivo* osteoclast differentiation assays treated with MERTK small molecule inhibitor R992 for 24h from culture day 3 to 4 (n=3, mean±SEM, unpaired t-test).

#### Supplemental Figure 4

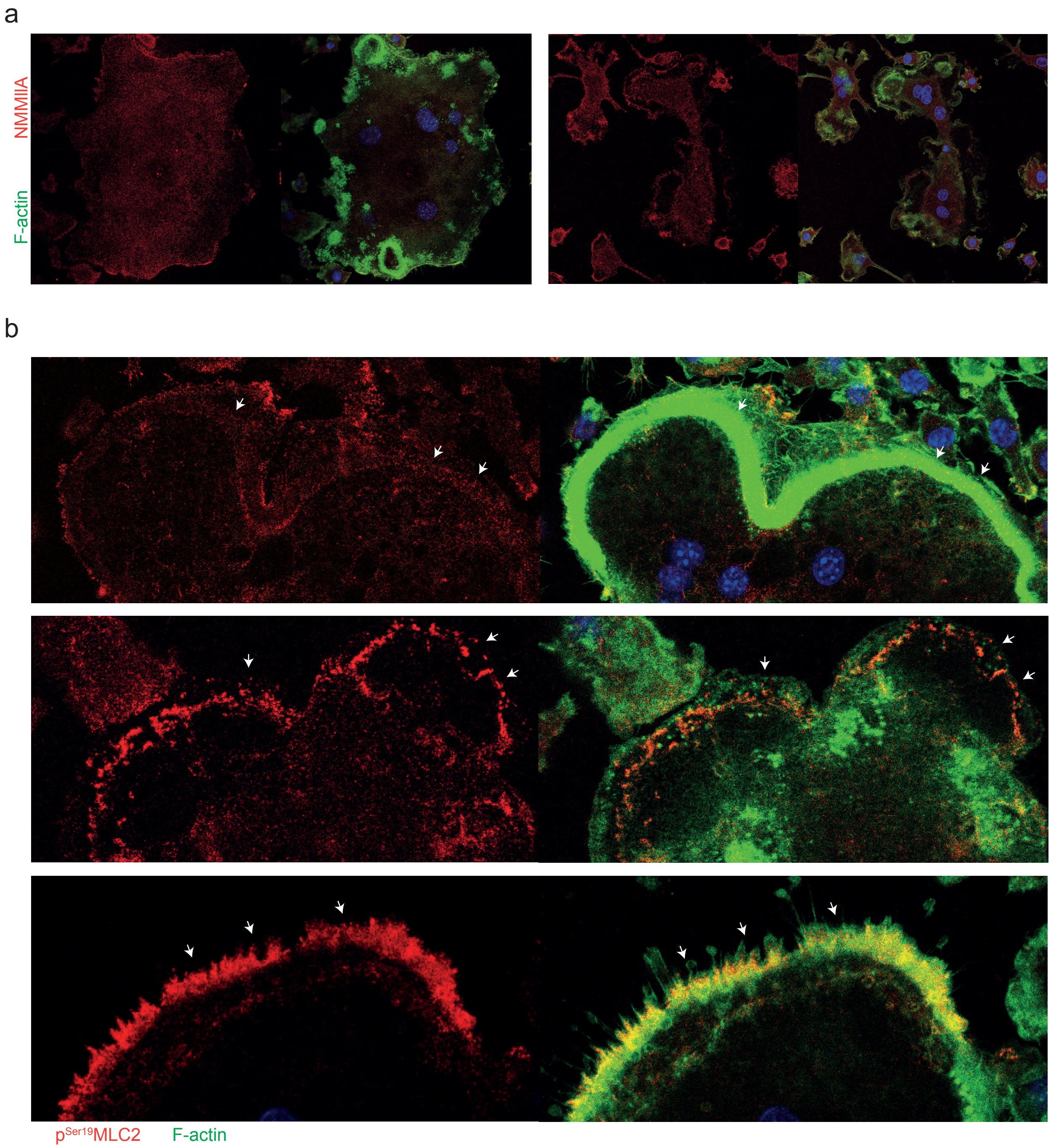


**Supplemental Figure 4. Spatial distribution of NMMIIA and p$^{Ser19}$MLC2 in osteoclasts. a,** Representative images of different osteoclast morphologies showing spatial distribution of NMMIIA in osteoclasts. **b,** Representative images of three different osteoclast morphologies showing spatial distribution of p$^{Ser19}$MLC2 in osteoclast adhesion structures and podosome belt.

### Supplemental Figure 5

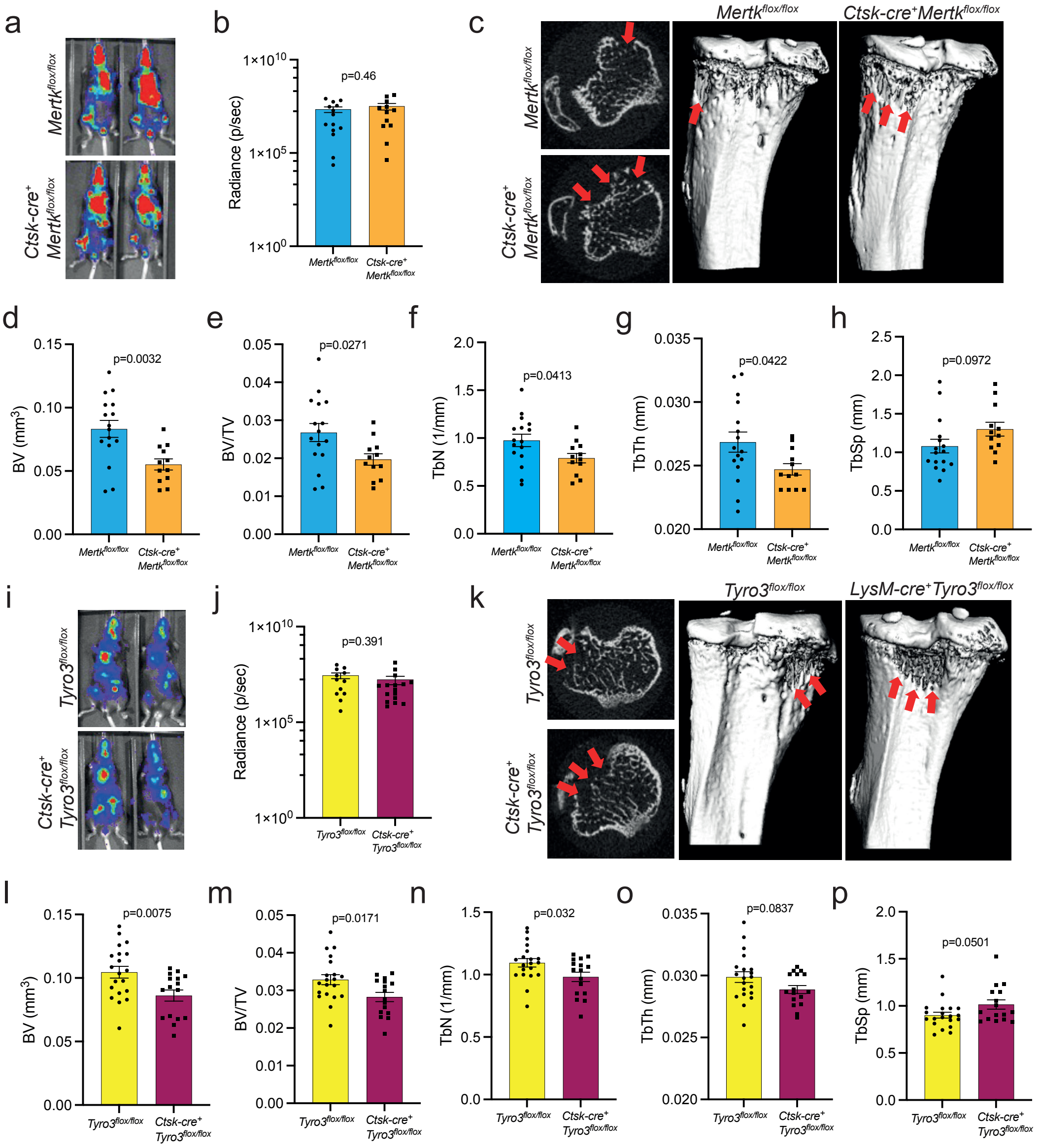

**Supplemental Figure 5. *cathepsin K*-mediated deletion of *Mertk* and *Tyro3* fosters osteolytic bone destruction of breast cancer bone metastases. a,** Luciferase⁺ EO771 breast cancer cells were injected intracardially in *Mertk$^{flox/flox}$* and *Ctsk-cre⁺Mertk$^{flox/flox}$* mice. Tumor spread was monitored via bioluminescence imaging (BLI). **b,** Analysis of tumor load in tibias by BLI after 10 days (n=14/12, mean±SEM, unpaired t-test). **c, d,e,f,g,h,** Representative µCT images and 3D reconstructions (c) and analysis (d-h) of BV, BV/TV, Tb.N, Tb.Th and Tb.Sp of trabecular bone of metaphyseal proximal region of the tibia of *Mertk$^{flox/flox}$* and *Ctsk-cre⁺Mertk$^{flox/flox}$* mice (n=16/12, mean±SEM, unpaired t-test). **i,** Luciferase⁺ EO771 breast cancer cells were injected intracardially in *Tyro3$^{flox/flox}$* and *Ctsk-cre⁺Tyro3$^{flox/flox}$* mice. Tumor spread was monitored via bioluminescence imaging (BLI). **j,** Analysis of tumor load in tibias by BLI after 10 days (n=16/12, mean±SEM, unpaired t-test). **k,l,m,n,o,p,** Representative µCT images and 3D reconstructions (k) and analysis (l-p) of BV, BV/TV, Tb.N, Tb.Th and Tb.Sp of trabecular bone of metaphyseal proximal region of the tibia of T*yro3$^{flox/flox}$* and *Ctsk-cre⁺Tyro3$^{flox/flox}$* mice (n=20/16, mean±SEM, unpaired t-test).
